# Molecular Mechanisms of Genotype-Dependent Lifespan Variation Mediated by Caloric Restriction: Insight from Wild Yeast Isolates

**DOI:** 10.1101/2024.03.17.585422

**Authors:** Samantha McLean, Mitchell Lee, Weiqiang Liu, Rohil Hameed, Vikas Anil Gujjala, Xuming Zhou, Matt Kaeberlein, Alaattin Kaya

## Abstract

Caloric restriction (CR) is known to extend lifespan across different species and holds great promise for preventing human age-onset pathologies. However, two major challenges exist. First, despite extensive research, the mechanisms of lifespan extension in response to CR remain elusive. Second, genetic differences causing variations in response to CR and genetic factors contributing to variability of CR response on lifespan are largely unknown. Here, we took advantage of natural genetic variation across 46 diploid wild yeast isolates of *Saccharomyces* species and the lifespan variation under CR conditions to uncover the molecular factors associated with CR response types. We identified genes and metabolic pathways differentially regulated in CR-responsive versus non-responsive strains. Our analysis revealed that altered mitochondrial function and activation of *GCN4-*mediated environmental stress response are inevitably linked to lifespan variation in response to CR and a unique mitochondrial metabolite might be utilized as a predictive marker for CR response rate. In sum, our data suggests that the effects of CR on longevity may not be universal, even among the closely related species or strains of a single species. Since mitochondrial-mediated signaling pathways are evolutionarily conserved, the dissection of related genetic pathways will be relevant to understanding the mechanism by which CR elicits its longevity effect.

**Author summary:** Caloric restriction **(CR)** is an energy-balanced nutrient intake without malnutrition to reduce food intake by 20-40%. CR leads to distinct metabolic reprogramming and adaptive changes in gene expression and, as a result, increases health and lifespan in various model organisms, from yeast to most likely primates. Besides extending lifespan, CR also holds great promise for treating many human age-onset pathologies, and the molecules underlying its effects are sought as targets of pharmaceutical aging therapeutics. However, despite extensive research, the mechanisms of lifespan extension in response to CR remain elusive. In addition, several studies in different aging models have now demonstrated that the longevity effect of CR can vary dramatically across different genotypes within a population. As such, CR might be beneficial for some yet detrimental for others, and the mechanisms underlying such genotype-dependent variation are not clear. In this study, we meet these challenges by dissecting molecular response to CR in diverse wild isolates of yeast strains, aiming to characterize pathways and molecules mediating CR’s effects on replicative lifespan (RLS) diversity. We found that the RLS significantly differs across genetically diverse wild yeast isolates under CR conditions. Examining the relationships among the RLS phenotypes under CR and non-CR conditions, transcript, and metabolite provided insights into the role of mitochondrial functions in CR-mediated lifespan extension.

## Introduction

Caloric restriction (**CR**) is an energy-balanced, non-invasive nutrient intake without malnutrition aimed at reducing food intake by 20-40% [1,2]. CR leads distinct metabolic reprogramming and adaptive changes in gene expression and, as a result, increases health and lifespan in various model organisms, from invertebrates to most likely primates, while also slowing down age-related diseases [1–5]. CR in yeast is modeled by simply reducing the glucose concentration in the growth medium from 2% to 0.05% (or less), which causes an increase in the replicative lifespan (**RLS**) [6–8], defined as the number of times a mother cell can divide [9]. Besides extending lifespan, CR also holds great promise for treating many human age-onset pathologies, and the molecules underlying its effects are sought as targets of pharmaceutical aging therapeutics [1–5, 10]. However, despite extensive research, the mechanisms of lifespan extension in response to CR remain elusive. In addition, several studies in different aging models have now demonstrated that the longevity effect of CR can vary dramatically across different genotypes within a population [7, 11–15]. As such, CR might be beneficial for some yet detrimental for others, and the mechanisms underlying such genotype-dependent variation are not clear [10]. A more integrated approach is needed to understand how the natural environment and natural selection interact to shape genotype and lifespan under CR condition.

In this study, we meet these challenges by dissecting molecular response to CR in ecologically and genetically diverse wild isolates of *Saccharomyces* species [16–18], aiming to characterize pathways that mediate the wide range of RLS phenotypes under CR conditions. We found that the longevity effect of CR varies dramatically within and between populations across different genotypes and species of budding yeast. Examination of the relationships between transcriptomes and the RLS phenotypes under CR and non-CR conditions provided insights into the mechanisms, mediated by *GCN4,* a nutrient-responsive transcription factor, and mitochondrial function-dependent CR effect on lifespan regulation, together, explaining a portion of heterogeneity in cellular processes that affect lifespan variation and CR responses.

Overall, we present evidence that mitochondrial-mediated mechanisms linking nutrition sensing to stress response pathways through *GCN4* is associated with the positive response to CR, and the abundance of mitochondrial metabolite n-formylmethionine can be utilized as a predictive marker for CR-mediated lifespan extension rate. Considering the conservation of pathways and the regulators, the principles of mechanisms learned through this work might apply to regulating lifespan in more complex organisms.

## Results

### The longevity effect of CR varies within and between populations across different genotypes and species of budding yeast

To analyze genotype-dependent responses to CR, we took advantage of natural genetic variation across 46 diploid wild yeast isolates of *Saccharomyces cerevisiae* (*S. cerevisiae*) and six other closely related budding yeast species; *S. kluyveri, S. bayanus, S. kudriavzevii, S. paradoxus, S. castellii, and S. mikatae.* We analyzed their RLS phenotype under both control (high glucose, 2%) and CR (low glucose, 0.05%) conditions.

Within *the S. cerevisiae* population, we found ∼10-fold median RLS variation under the control condition, and RLS analysis under the CR condition revealed a 7-fold variation among them (**Figure 1A**, Supplementary File S1). While 26 strains displayed significant (Wilcoxon rank sum test, adjusted *P* ≤ 0.05) CR response (either decreased or increased) in median RLS, the remaining 20 strains did not respond to CR (no significant changes in median RLS) (**Figure 1A**, Supplementary File S1). Among the 26 CR responding strains, 11 of them showed various degrees of median RLS extension, while the remaining 15 displayed a wide range of decreased median RLS (**Figure 1A**, Supplementary File S1). For example, under the CR condition YJM978 strain showed a decreased median RLS (50% decrease), whereas CR caused a 75% median RLS increase of the Y9 strain compared to the control condition (**Figure 1B**, Supplementary File S1). Consistent with the previous report [8], the control diploid laboratory WT strains BY4743, which is a derivative of the original S288c isolate, also increased median RLS by 12% (Wilcoxon rank sum test, adjusted *P* = 0.009). On the other hand, the original S288c strain did not respond to CR (**Figure 1B**), indicating adaptive changes to laboratory conditions resulted in an alteration to CR response in this strain background.

**Figure. 1:**
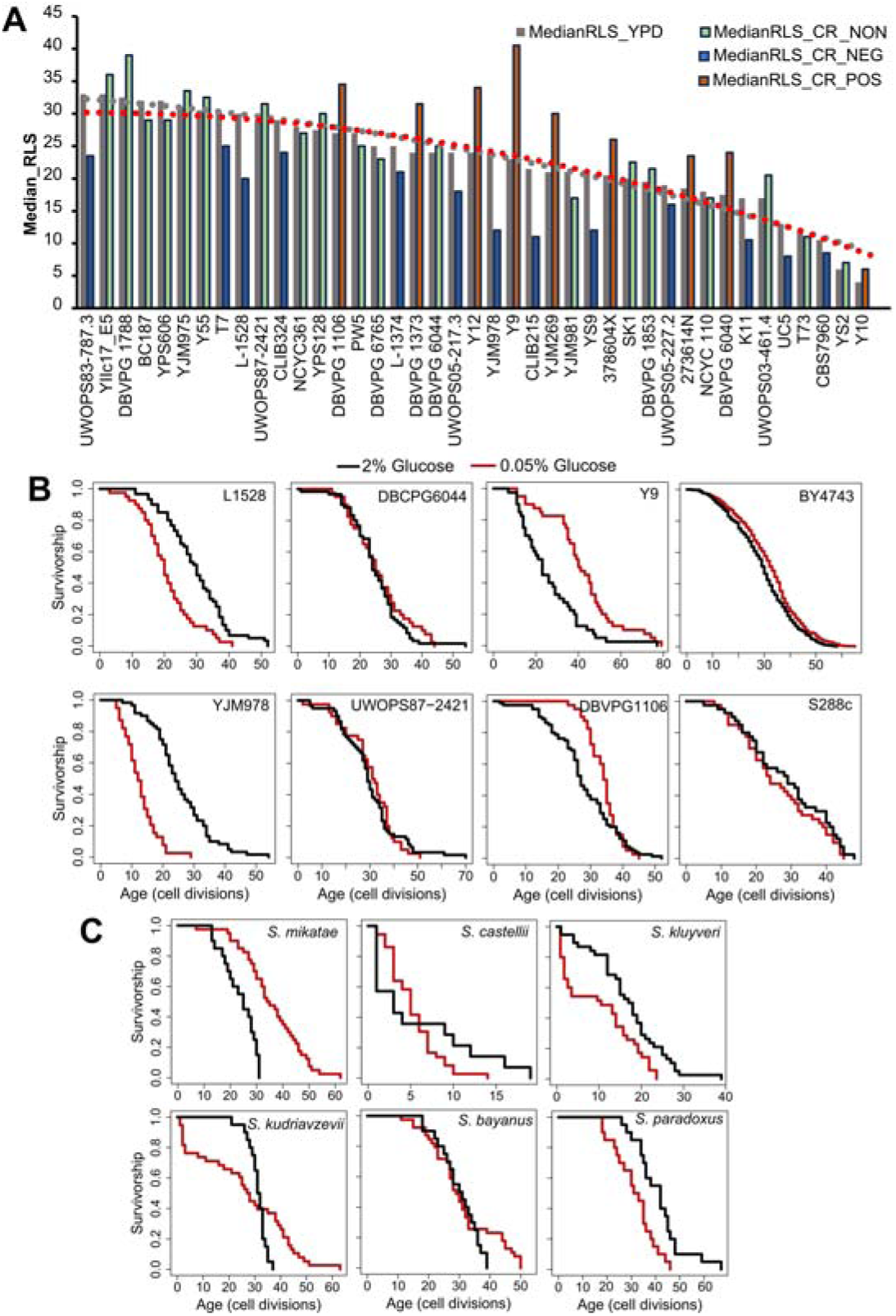
Intra- and interspecies distribution of median RLS distribution. (**A**) Cells were grown in yeast peptone dextrose (YPD). Dashed lines represent the average median RLS of YPD (2% glucose gray) and YPD-CR (0.05% glucose). The bar graph depicts the media RLS of each strain analyzed under YPD (gray) and YPD-CR. Bar colors indicate CR response types based on the statistical significance of median RLS changes under CR conditions compared to the YPD. Blue is for negatively responding strains (NEG), green is for non-responding strains (NON), and orange bars represent positively responding strains (POS). (**B**) Examples of lifespan curves for the selected strains of *S. cerevisiae*. (**C**) Lifespan curves for six *Saccharomyces* species. The black curve shows the lifespan under YPD conditions, and the red curve shows the lifespan under YPD CR conditions. The raw data and statistical significance can be found in Supplementary File 1.

Next, to understand whether the longevity effect of CR also varies across different species, we also analyzed the RLS of six different budding yeast species of *Saccharomyces* genus; *S. kluyveri, S. bayanus, S. kudriavzevii, S. paradoxus, S. castellii, and S. mikatae*. Interestingly, only two of them, *S. mikatae* and *S. kluyveri* showed significant (Wilcoxon rank sum test, adjusted *P* ≤ 0.05) lifespan (median RLS) extension when subjected to the CR (**Figure 1C**, Supplementary File S1).

Overall, this data showed considerable variation in lifespan phenotype under CR conditions among the genetically diverse natural isolates of the same species and the closely related species of the same genus. This data suggests that CR at the 0.05% glucose level does not promote lifespan extension for most of the wild-derived yeast strains and species, and this applies to other non-laboratory adapted model organisms.

### Comparison of gene expression pattern between CR responding and non-responding strains

Gene expression variation has been suggested to play a significant role in adaptive evolution. Existing research also highlights the potential influence of gene expression levels on various phenotypic traits and its plasticity [19,20], such as changes in lifespan [21,22].

Accordingly, we examined whether the comparison of transcriptome profiles of these strains, obtained under the high glucose condition, can reveal molecular signatures that can predict the CR response type. First, we examined whether evolutionary relationships based on the gene expression variation are associated with the CR response types; we constructed gene expression phylograms for *S. cerevisiae* strains using a distance matrix of 1 minus Spearman correlation coefficients [23] and based on normalized reads [16]. We found that CR response types are mainly branched through early adaptation of strain-specific life history trajectories at gene expression level, and this adaptation is mostly acquired independently (**Figure 2A**). Subsequently, principal component analysis (PCA) was conducted, and the result did not reveal a distinct segregation pattern, except for a few outlier strains that were separated by PC1 and PC2. Intriguingly, this separation resembled the phylogenetic relationship, with the cumulative effect of the first three principal components accounting for approximately 35% of the total variance in gene expression (**Figure 2B**). We then employed a different dimensionality reduction method, partial least squares discriminant analysis (PLS-DA), particularly suited for distinguishing different groups (Please see methods). PLS-DA analyses revealed distinct clusters for positively responding strains (**Figure 2C**). The model evaluation metrics provided further validation, with an R2Y value of 0.926 indicating that the model accounts for 92.6% of the variability in these three groups and a Q2Y value of 0.766, suggesting a predictive accuracy of 76.6%.

**Figure 2:**
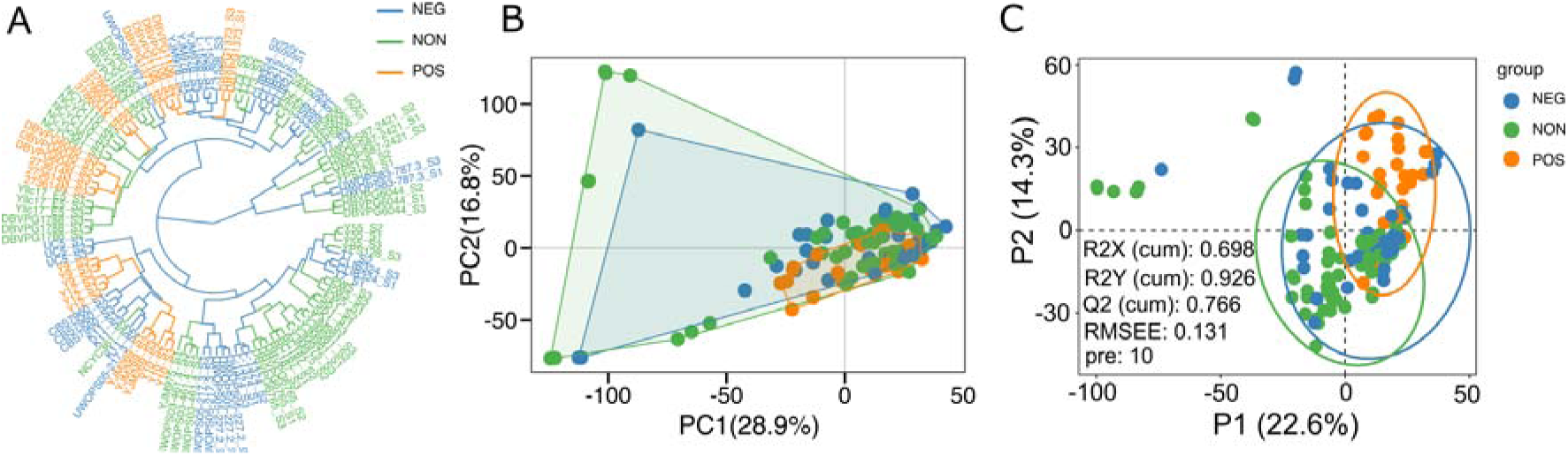
Sample clusters. (**A**) Dendrogram plot of sample cluster. Clustering is generated by hierarchical clustering using the Ward D2 method. The color represents three groups, as shown in Figure 1; Orange represents the positively responding (POS) group. Green represents the non-responding (NON) group. Blue represents the negatively responding (NEG) group. (**B**) Principal component analysis of gene expression across three groups. The first two principal components (PCs) and their variance explanation percentages are shown. Each repetition is treated as a point. (C) Partial least squares discriminant analysis (PLS-DA) based on the gene expression data across wild isolates of three phenotypic groups is shown. The scatter plot of the first two partial least squares (PLS) components and their variance explanation percentages are shown. The model parameters are also shown in the figure, including the explanatory degree of the model to independent variables (R2X), the explanatory degree of the model to dependent variables (R2Y), the predictive ability of the model (Q2Y), the root mean square error (RMSEE) and the number of PLS components used when calculating these parameters (pre). Each sample is treated as a point. Each phenotypic group is represented by different colors.

Next, to identify molecular signatures and genetic regulators of CR response types, we compared the gene expression pattern between CR responding strains (increased median RLS) to non-responding strains (no significant change) and negative responding strains (decreased median RLS). Under high glucose conditions, 222 genes differentially expressed (DEGs), including 146 genes that had significantly reduced expression and 76 genes that had increased expression in the CR responding group in comparison to the non-responding group (adjusted *P* ≤ 0.05 and log2-fold > 0.5) (**Figure 3A, Supplementary File S2)**. Similar analysis revealed 176 DEGs, including 50 genes with significantly reduced expression and 120 genes with increased expression in the CR responding group compared to the negative-responding group (**Figure 3B, Supplementary File S2)**.

**Figure 3:**
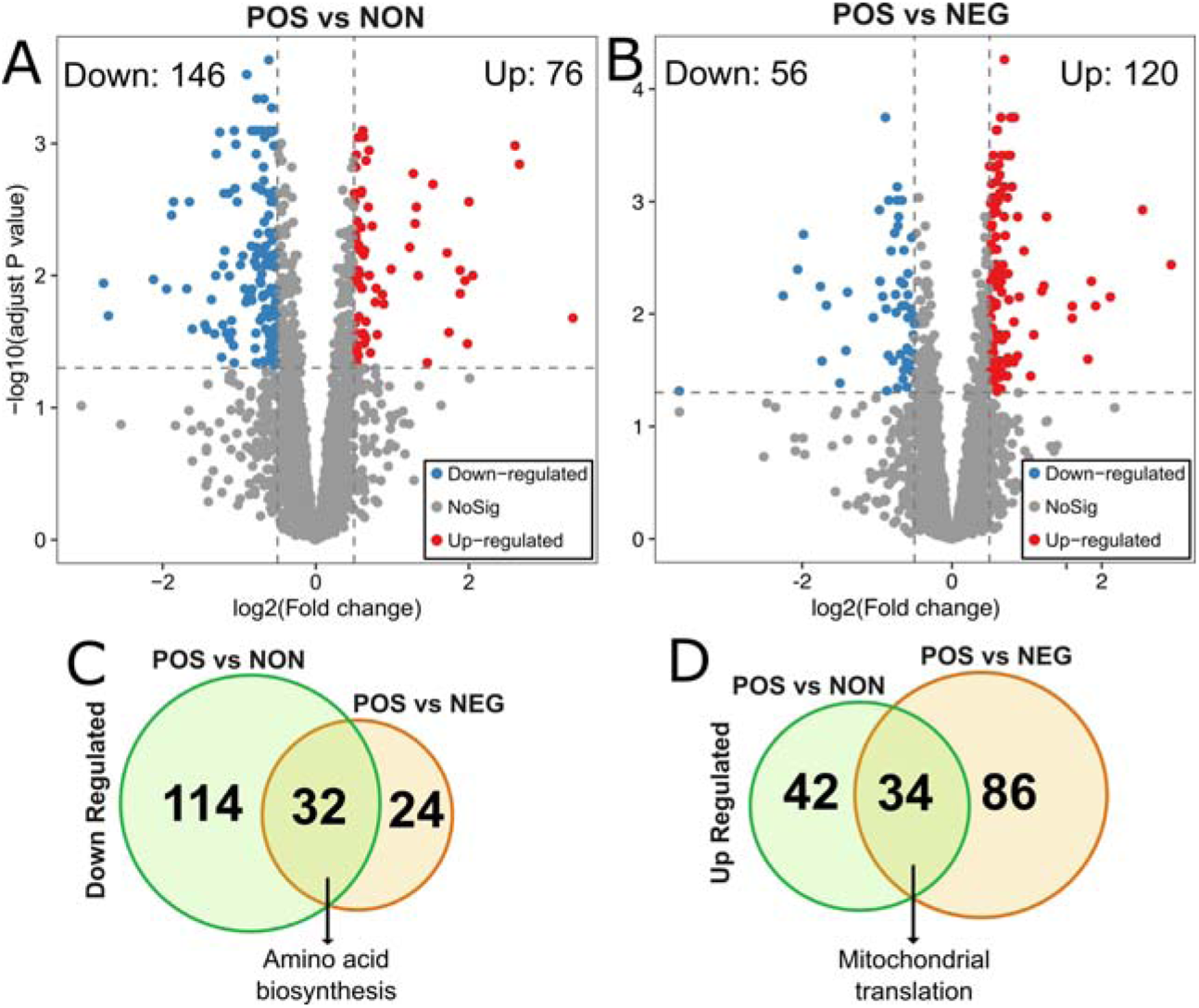
Differentially expressed genes in POS group compared to the NON and NEG groups. The volcano plots depicting differentially expressed genes (DEGs) resulted in positively responding (POS) versus (**A**) non-responding (NON) groups and (**B**) negativelu-responding (NEG) groups. Red dots represent genes expressed at higher levels, while blue dots represent genes with lower expression levels in the POS group for each comparison. Gray dots represent genes that do not show significant differential expression between both groups. The gray line represents the cutoff of significant differential expression (adjust P value < 0.05 and log2 (Fold change) >0.5). The Y-axis denotes −log10 (adjusted *P* value) while the X-axis shows log2 (Fold change). Venn diagram shows the number of unique and common DEGs from each comparison (**C**) for down-regulated and (**D**) for up-regulated DEGs. The complete list of DEGs with p values can be found in Supplementary File 2.

To further investigate the molecular patterns associated with the CR response type, we analyzed the commonalities and differences in gene expression between two groups (positively responding vs non-responding and positively responding vs negatively responding). Among the down-regulated DEGs identified from both comparisons, 32 genes were in common. The GO term associated with these commonly down-regulated genes in positively responding strains was enriched in cellular amino acid biosynthesis (**Figure 3C**). Among the up-regulated DEGs, there were 34 genes in common. Functional enrichments of these commonly up-regulated DEGs resulted in a single GO term associated with mitochondrial translation (**Figure 3D**).

We then performed GO enrichment and KEGG pathway analysis for the uniquely altered genes in positively responding strains compared to the non-responding or negatively responding strains alone. The 114 unique down-regulated DEGs, resulting from the comparison of positively responding versus non-responding strains, revealed that positively responding strains have decreased lipid and amino acid biosynthesis and metabolism compared to the non-responding strains. (**Figure 4A, B)**. A similar result was also obtained for 24 down-regulated DEGs from positive versus negative responding strains comparison (**Figure 4C, D)**. The up-regulated 42 unique DEGs resulted from a comparison of positively responding versus non-responding strains enriched in the ribosome (both mitochondrial and cytosolic) and mRNA surveillance pathways. Additionally, GO term enrichment indicated an increase in iron metabolism in positively responding strains (**Figure 4A, B)**. The GO terms for 86 up-regulated DEGs resulted from positively responding versus negatively responding strains comparison were enriched in mitochondrial translation, ATP synthesis coupled proton transport, ATP biosynthetic process, and mitochondrial respiratory chain complex IV assembly (**Figure 4C, D**). Overall, these results suggest that responding strains maintain higher translation under high glucose conditions and are uniquely adapted to regulate mitochondrial function for energy production compared to the non-responding and negatively responding strains. In addition, our data suggests that negative responding strains are characterized by high amino acid biosynthesis and metabolisms. On the other hand, non-responding strains might also be compromised for iron and copper homeostasis. Further analyses of KEGG pathways and GO term enrichment from each comparison revealed decreased fatty acid and branched-chain amino acid synthesis in positively responding strains. At the same time, these two processes are up-regulated in other groups under high glucose conditions (**Figure 4C, D, and Supplementary Figure S1)**. Overall, we showed a phenotype-specific gene expression that, in comparison to the non- or negatively responding strains, specific group genes were selectively up or down-regulated in positively responding strains under high glucose conditions (**Figure 5 and Supplementary File S2**).

**Figure 4:**
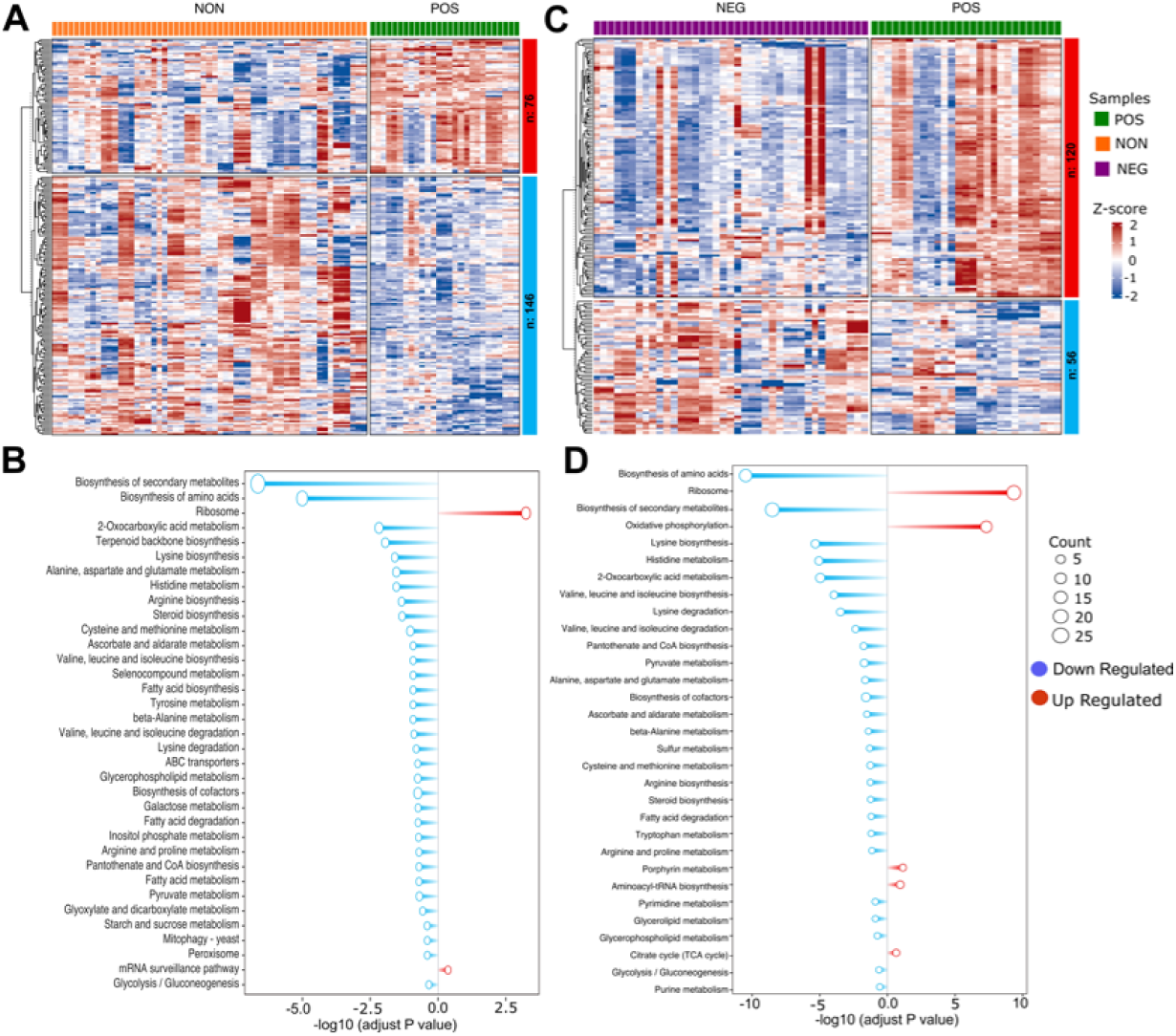
Gene enrichment analysis of different expressed genes. (**A**) Heatmap depicting differentially expressed genes (DEGs) in positively responding (POS) versus non-responding (NON) groups. Each column represents an individual strain, and each row represents a single gene expression change between the POS and NON groups. The color intensity indicates the level of gene expression, and the expression value is scaled (Z-score). Red indicates higher expression, and blue indicates lower expression. (**B**) Bar plot depicting gene enrichment analysis of DEGs. The Y-axis shows each significantly enriched KEGG pathway while the X-axis denotes −log10 (adjusted *P* value). Red bars represent pathways expressed at higher levels in the POS group, while blue bars represent pathways with higher expression levels enriched in the NON group. Point size represents the number of genes in this pathway. (**C and D**) Similar analyses were also done for POS and NEG groups. The complete list of DEGs with p values can be found in Supplementary File 2.

**Figure 5:**
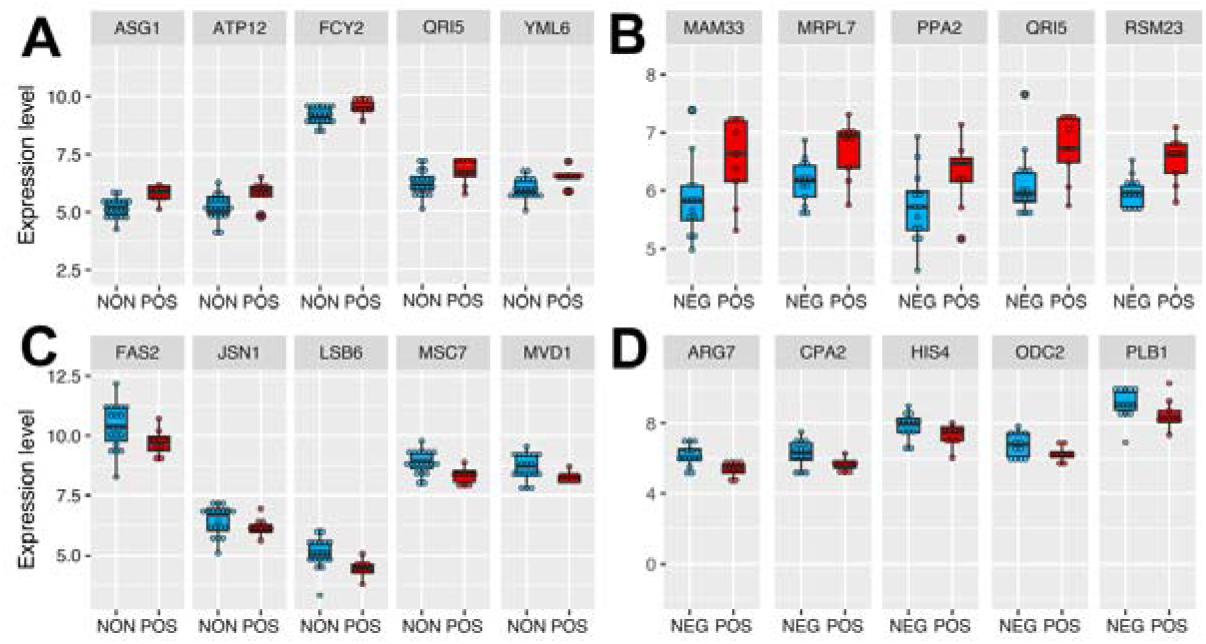
Top differentially expressed genes in responding strains from each comparison. The box plot depicts the top genes significantly up-regulated in the positively responding (POS) group compared to the **(A)** non-responding (NON) and (**B**) negatively responding (NEG) groups. (**C and D**) The lower panels contain examples of significantly down-regulated genes in the POS group resulting from each comparison. The Y-axis denotes gene expression level, and the X-axis shows two groups. Expression values across samples can be found in Supplementary File 2.

Next, to further understand the relationship between gene expression variation and CR-mediated lifespan variation, we performed, phylogenetic regression by generalized least squares (PGLS), as we described previously [16,17]. Our analyses further showed that expression levels of some of these genes were significantly correlated with CR response variability across all strains (**Figure 6 and Supplementary File S2**). For example, we identified 118 transcripts with significant correlation with median RLS (adjusted *P* ≤ 0.01; 48 with positive correlation and 70 with negative correlation) (**Supplementary File S2**). Among the top hits with positive correlation were a major mitochondrial D-lactate dehydrogenase (*DLD1*-oxidizes D-lactate to pyruvate), homeodomain-containing protein and putative transcription factor (*TOS8*), and SKI complex-associated protein (*SKA1*-involved in involved in 3’-5’ degradation of long 3’UTR-containing mRNA) (**Figure 6A**). The top hits with negative correlation included the genes coding for DNA end-binding protein required for nonhomologous end joining (*NEJ1*), GTPase-activating protein for Gpa1p (*SST2*), and Subunit of G protein involved in pheromone response (*GPA1*) (**Figure 6A**). The data indicates a role for *GPA1* function in regulating lifespan under CR condition, which plays a role in mating-related nuclear migration and karyogamy also involved in inositol lipid-mediated signaling and regulation of MAPK export from the nucleus.

**Figure 6:**
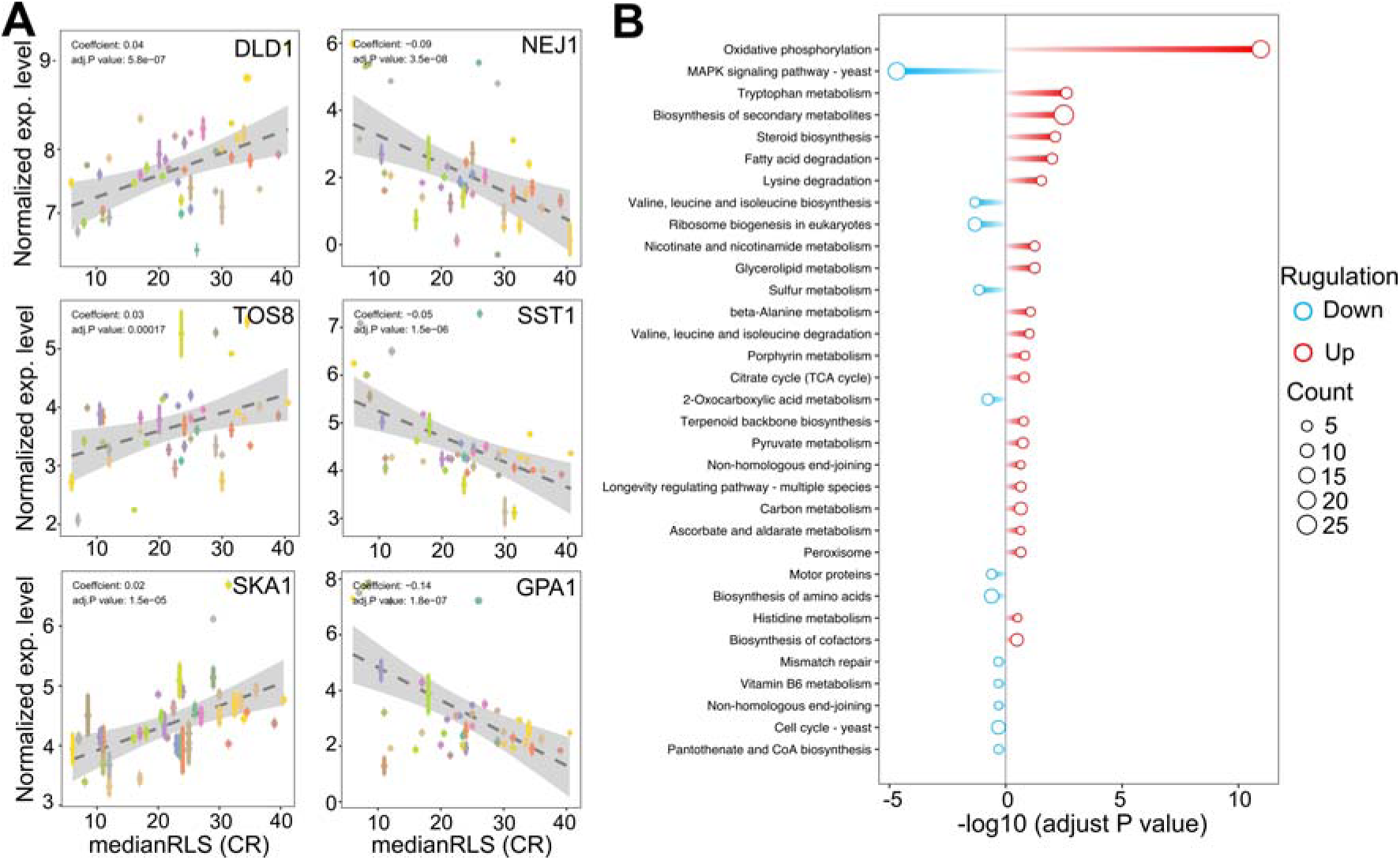
Selected genes whose expression under high glucose condition correlates with median replicative lifespan (RLS) under CR condition. (**A**) *DLD1*, *TOS8*, and *SKA1* gene expression levels, determined under high glucose condition correlate positively, and *NEJ1, SST1*, and *GPA1* correlate negatively with median RLS under CR conditions. The Y-axis denotes the expression level, and the X-axis represents the lifespan. Each color point represents an individual strain. Error bars represent standard error (SE). The gray area represents a 95% confidence interval. The regression coefficient and adjusted *P* values are included in the figure. The complete list of significantly correlating genes, regression slopes, and p values can be found in. Supplementary File 2. (**B**) Bar plot depicting gene enrichment analysis of lifespan-correlated genes. The Y-axis shows each significantly enriched pathway of the KEGG database while the X-axis denotes −log10 (adjusted *P* value). Red bars represent pathways expressed at higher levels in higher lifespan strains, while blue bars represent pathways with higher expression levels in lower lifespan strains. Point size represents the number of genes in this pathway. The complete list of genes with regression slopes, and p values can be found in Supplementary File 2.

In agreement with the cross-comparison transcriptome data, the KEGG pathway analysis of these positively correlated genes revealed that strains with increased oxidative phosphorylation activity and amino acid and fatty acid degradation pathways show effective CR response (**Figure 6B**). Contrarily, negatively correlating genes enriched in MAPK signaling pathway, branched-chain amino acid biosynthesis, mismatched repair, and cell cycle (**Figure 6B**).

In conclusion, the comprehensively analyzed transcriptome data across diverse genetic backgrounds and the lifespan variation under CR conditions uncover the molecular factors associated with CR response types.

### Comparison of metabolite abundance pattern between CR responding and non-responding strains

We also searched for metabolites whose abundances identified under high glucose condition are associated with CR response type. The metabolome represents a snapshot of relevant biological processes downstream of the proteome, and it has been widely used for characterizing aging-regulated metabolic pathways and biomarkers for age-associated diseases [24–26]. Among the 166 metabolites that we examined, none of them showed significant differences at adjusted p-value cut-off (p ≤ 0.05) in both responding versus non-responding and responding versus negatively responding comparisons (**Supplementary File S2**). At the non-adjusted p-value cut-off (p ≤ 0.01), we found responding strains with decreased abundance of leucine and S-Adenosyl-homocysteine and increased abundance of inositol monophosphate in comparison to both non- and negatively responding groups. Three metabolites, nicotinic acid, mevalonate, and 1-metyhylnicotinamide were specifically characterized by decreased abundance in responding strains in comparison to the non-responding group. Oxidized glutathione and phenylalanine were specifically characterized by decreased abundance in positively responding strains in comparison to the negatively responding group (**Figure 7A, B and Supplementary File S2**).

**Figure 7:**
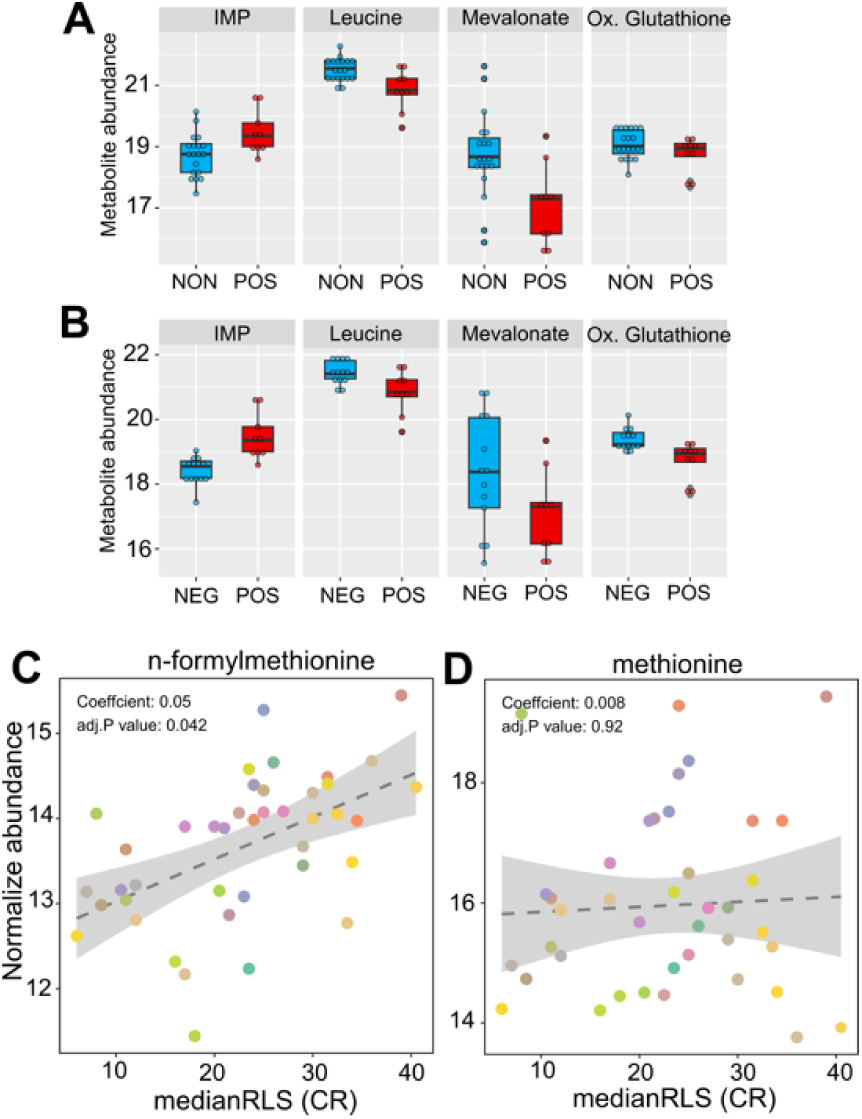
High glucose metabolite abundance patterns are associated with median RLS under CR condition. The box plot depicts the top metabolites significantly altered in the positively responding (POS) group, compared to the **(A)** non-responding (NON) and (**B**) negatively responding (NEG) groups. (**C**) Positive significant correlation of n-formylmethionine and (**D**) non-significant correlation of methionine with median RLS. The Y-axis denotes the abundance level of each metabolite identified under high glucose condition, and the X-axis represents the median RLS, determined under CR condition. Each color point represents an individual strain. The gray area represents a 95% confidence interval. The complete list of metabolites with normalized abundance or regression slopes, and p values can be found in Supplementary File 2.

Next, we examined the correlation between metabolite abundance and CR-mediated lifespan variations. Our phylogenetic regression (PGLS) analyses identified a single metabolite, n-formylmethionine (fMet), with a significant (adjusted *P* = 0.041) positive correlation (**Figure 7C, Supplementary File S2**). Methionine (Met) derivate fMet initiates protein synthesis in mitochondria. Our data showed that fMet abundance was not coupled to Met abundance since there was no significant correlation between Met and median RLS (**Figure 7D, Supplementary File S2**). Given that both transcriptomic and metabolomic data were obtained under high glucose conditions, our regression analysis was not aimed at predicting CR response type. However, it gives an association between CR-mediated lifespan extension rates and molecule abundance which can be utilized as a predictive marker.

### CR mediates response type specific molecular changes

Next, to further investigate CR-mediated strain-specific transcriptional changes, we selected two CR responding (DBVPG1106, DBVPG1373) and non-responding (BC187, DBVPG6765) strains and collected cells that were grown under YPD-CR condition and subjected them transcriptomics analyses. We observed a distinct transcriptional response between responsive and non-responsive strains under CR conditions (**Figure 8A, B**). Our calculation of distances between strains based on the biological coefficient of variation (BCV) [27] of gene expression revealed two separate clusters. The CR regimen caused transcriptional changes that segregated the non-responding strains, mainly BCV1 and BCV2. Conversely, CR responding strains only separated along BCV2 (**Figure 8B**). The segregation pattern indicates unique transcriptional changes among the responding and non-responding groups. To analyze DEGs, we analyzed CR-mediated changes compared to the controls (high glucose) for each group. We found that CR caused more robust expression changes in the non-responding group. Overall, there were 2185 DEGs (1213 down-regulated and 972 up-regulated) significantly altered (adjusted *P* ≤ 0.01, log2-fold change ≥ 1) in the non-responding group when subjected to CR. On the other hand, we identified only 422 DEGs (230 down-regulated and 192 up-regulated) significantly altered in the positively responding group when subjected to CR (**Figure 8C, Supplementary File S3**). Among them, 134 down-regulated and 110 up-regulated genes were shared between the responding and non-responding groups (**Figure 8C, Supplementary File S3**). These commonly altered genes are enriched in cytoplasmic translation for down-regulated and Gluconeogenesis, TCA cycle, Biosynthesis of nucleotide sugars, Tryptophan metabolisms and Longevity regulating pathways, and Fatty Acid degradation for up-regulated genes (**Supplementary File S3**).

**Figure 8:**
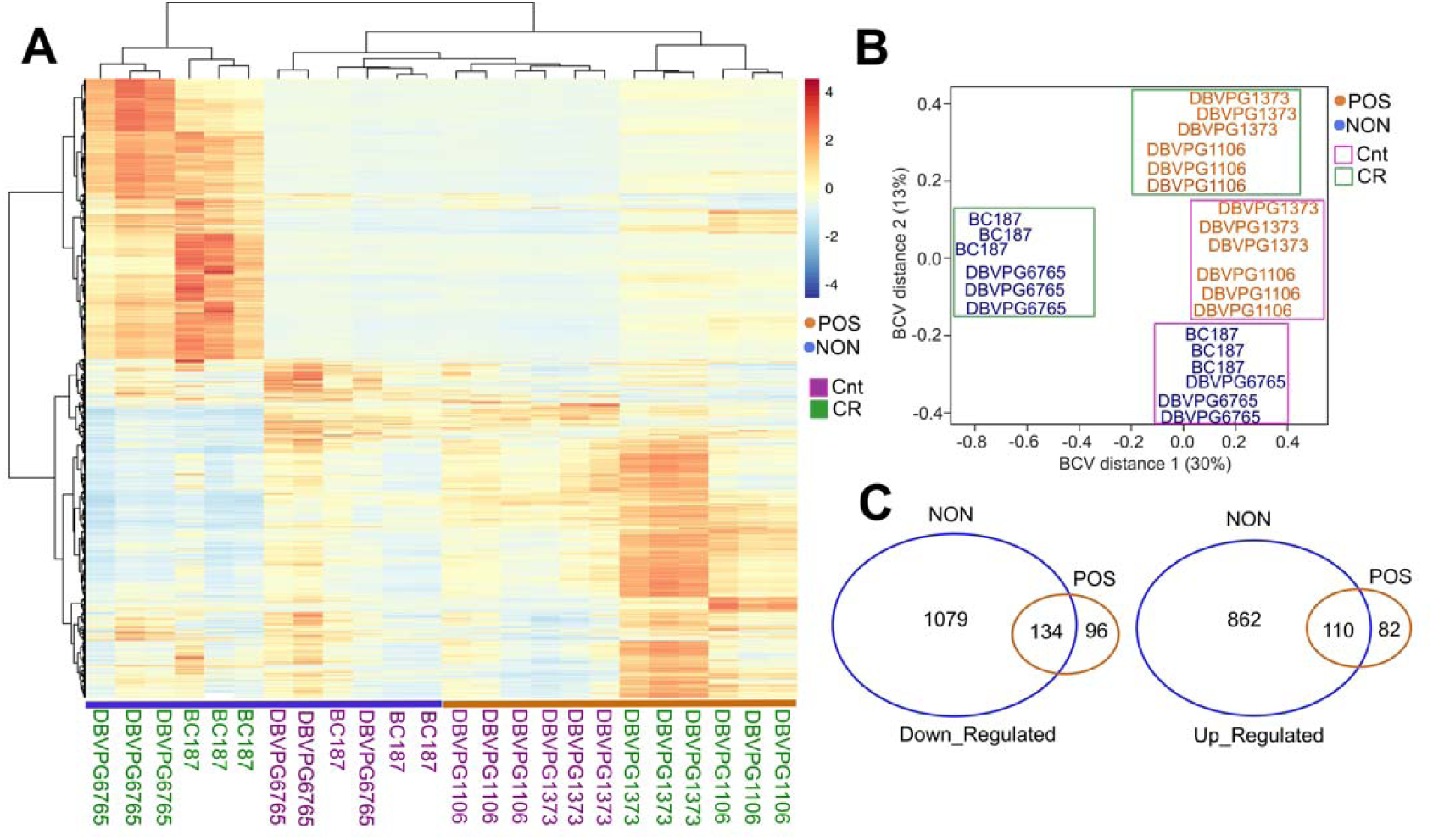
CR mediates phenotype specific transcriptional changes in responding and non-responding strains. **(A)** Heat map shows the differentially expressed genes (DEGs) and clusters for selected responsive (POS; orange) and non-responsive (NON; blue) strains under CR conditions in comparison to the matching control groups. (**B**) Calculation of distances between strains based on the biological coefficient of variation (BCV) of CR-mediated gene expression revealed two separate clusters for responding and non-responding strains. (**C**) Number of common and unique DEGs for each group. The left diagram shows down- and the right diagram shows up-regulated DEGs. The raw and normalized reads as well as DEGs with log2-fold change values along with statistical significance can be found in Supplementary File 3.

Then, we analyzed response type-specific DEGs for each group. For the non-responding group, we found that CR induces gene expression changes that up-regulate the pathways associated with Carbon metabolism, TCA cycle, Oxidative phosphorylation, Propanoate metabolisms, Branched Chain amino acid and Lysine degradation, Longevity regulating pathway, Biosynthesis of cofactors, and Autophagy (**Figure 9A**). Among the down-regulated pathways were Ribosome biogenesis, Amino acid biosynthesis and metabolism, and Nucleotide metabolisms (**Figure 9B**). Analysis of DEGs in CR responding strains revealed that CR caused down-regulation of Ribosomal genes and Tryptophan metabolisms and upregulation of MAPK signaling, TCA cycle, Amino acid metabolic pathways, and Biosynthesis of secondary metabolites. This data suggests that CR selectively acts on specific sets of genes to alter targeted pathways to extend lifespan in responding strains (**Figure 10A, B)**. For example, responding strains are characterized by higher mitochondrial respiration and translation under high glucose conditions (**Figure 4B, D**). Accordingly, the altered genes were only related to decreased ribosomal biogenesis, and there were no further alterations for genes associated with mitochondrial function in responding strains under CR conditions. However, CR-induced expression of genes alters mitochondrial function and decreases ribosomal biogenesis and translation for non-responding strains. Considering that the non-responding strains are characterized by decreased ribosomal biogenesis and translation under high glucose conditions (**Figure 4B, D**), nonselective repression of ribosomal biogenesis genes under CR conditions might further decrease translation. Thus, the imbalance between mitochondrial and cytosolic translation might be a confounding factor for preventing CR-mediated lifespan extension in non-responding strains, as suggested previously [28].

**Figure 9:**
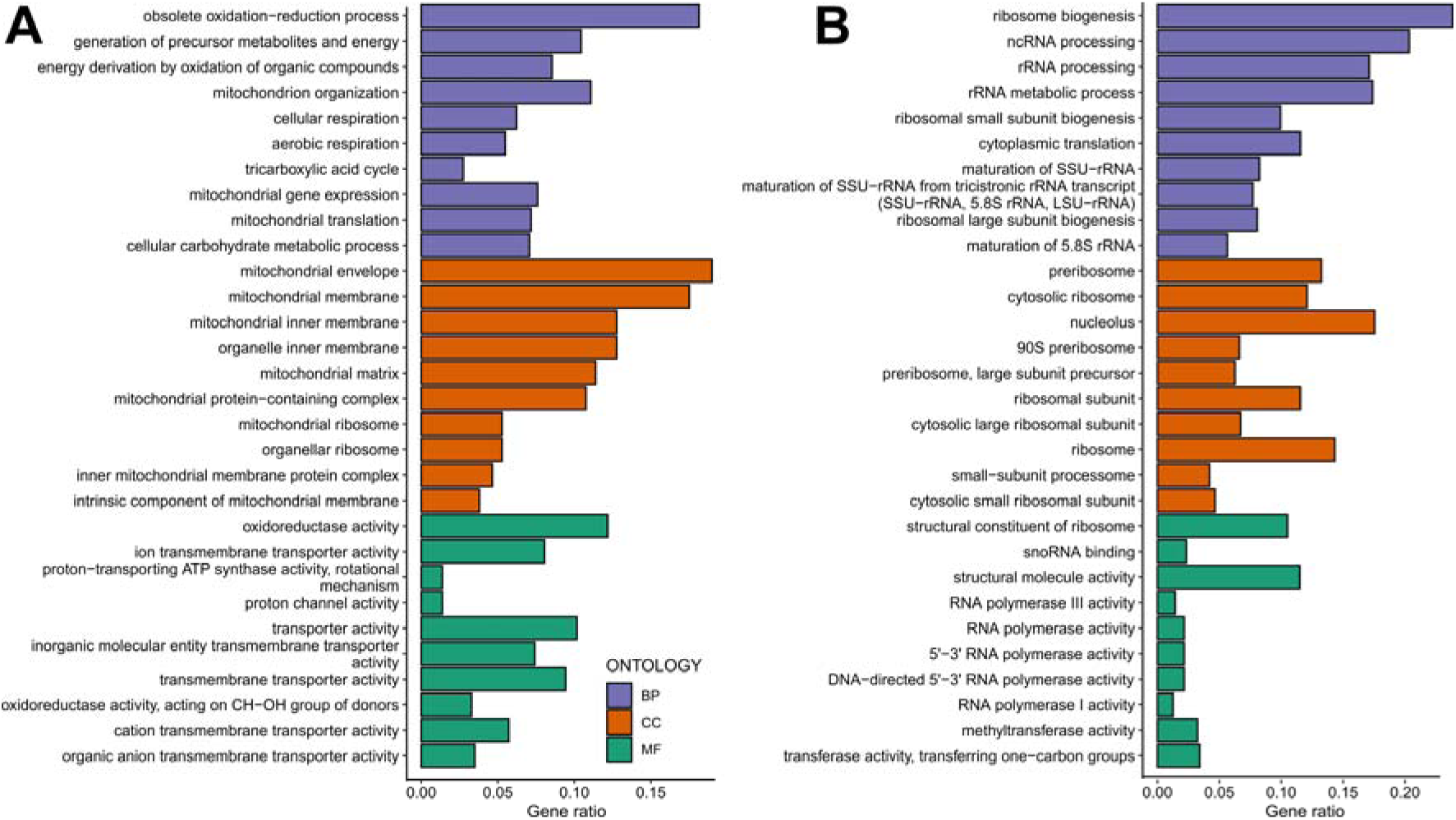
Gene ontology enrichment analyses mediated by CR in non-responding strains. Gene ontology enrichment analyses for the response type-specific **(A)** up-regulated DEGs and (**B**) down-regulated DEGs for the non-responding strains. GO categories include Biological Process (BP-Purple), Cellular Component (CC-Orange), and Molecular Function (MF-Green). The raw values of enrichment analyses, statistical significance, and genes associated with each term can be found in Supplementary File 2.

**Figure 10:**
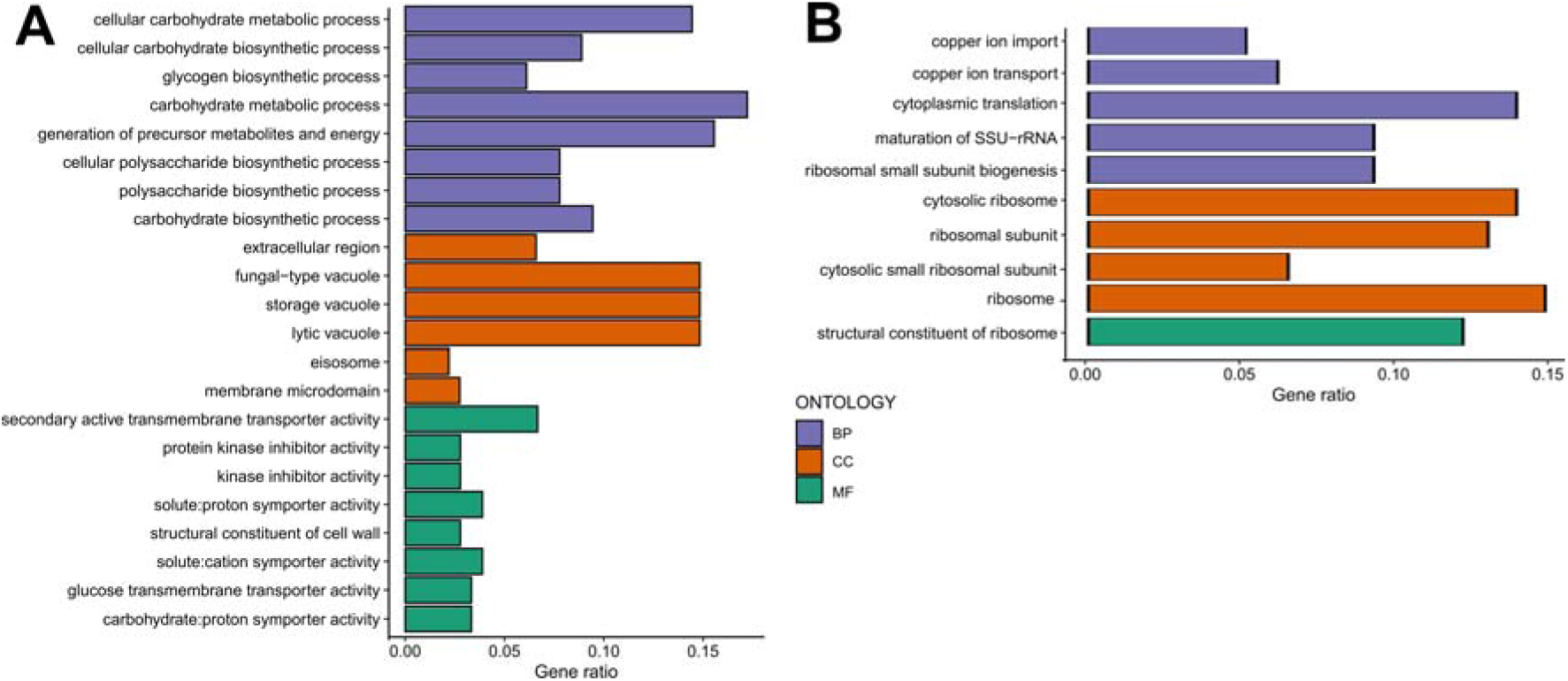
Gene ontology enrichment analyses mediated by CR in responding strains. Gene ontology enrichment analyses for the response type-specific **(A)** up-regulated DEGs and (**B**) down-regulated DEGs for the non-responding strains. GO categories include Biological Process (BP-Purple), Cellular Component (CC-Orange), and Molecular Function (MF-Green). The raw values of enrichment analyses, statistical significance, and genes associated with each term can be found in Supplementary File 2.

Next, to understand the regulatory network under CR conditions, we identified core Transcription Factors (TF) for up- and down-regulated genes of responding and non-responding strains. In responding strains, increased expressions of genes are regulated by several transcription factors, including *SKN7, RPN4, GCN4, HSF1, MSN2/4, YAP1,* and *MIG1* (**Figure 11A**). Many of these transcription factors regulated genes associated with environmental stress response genes (ESR) [29]. In fact, 53 of the 192 up-regulated genes in responding strains belonged to the ESR group (**Supplementary File S3**). Furthermore, our data suggests that many of the ESR genes and other up-regulated genes are regulated by *GCN4* (**Figure 11B**). On the other hand, decreased expressions of genes are regulated by *HMS1, CRZ1,* and *UME1* (**Figure 11C**) in the responding strains.

**Figure 11.**
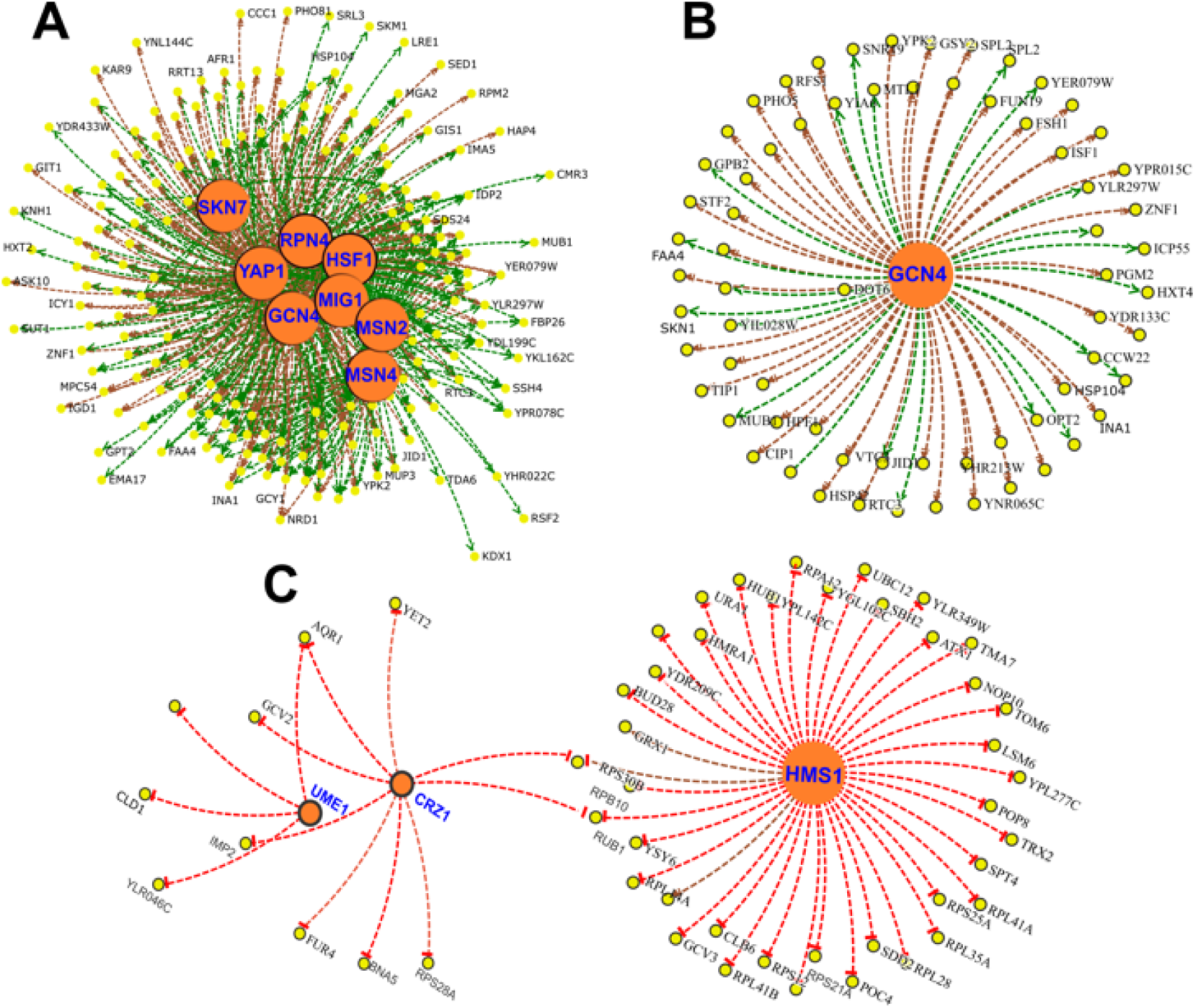
Gene regulatory network of DEGs altered under CR condition in responsive strains. Transcription Factors (TFs) (**A**) associated with up-regulated DEGs and (**B**) subnetwork of *GCN4* targets. (**C**) TF network for down regulated genes. Yellow nodes indicate gene targets that are altered in responsive strain under CR condition. Orange nodes indicate corresponding TFs. Green lines indicate gene activation and red lines represent transcriptional repression.

For the nonresponding strains, the increased expressions of genes were regulated by *MET32, MSN2, HCM1, HAP4, AFT2,* and *AFT1* under CR condition (**Supplementary Figure S2A**). The data suggests that increased expressions of respiratory genes were regulated by *HAP4* and *HCM1* in non-responding strains. *ATF1* and *ATF2* regulate iron homeostasis, while *MET32* is involved in transcriptional regulation of the methionine biosynthetic and other sulfur metabolic genes. Among the top TFs associated with down-regulated genes were *SFP1, CST6, HSF1, RAP1,* and *FHL1* (**Supplementary Figure S2B)**. Many of these TFs are involved in the regulation of ribosomal gene expression and might be associated with decreased translation in non-responding strains.

### Experimental testing of mitochondrial role and GCN4 function in CR-mediated lifespan extension

We further dissected the effect of mitochondrial respiration on CR response by comparing median RLS variation between YPD-CR and YPG (3% glycerol) conditions. A respiratory growth substrate, glycerol can extend RLS [30,31] by a switch from fermentation to respiration; however, the exact mechanism of glycerol-mediated lifespan extension is unclear. We previously observed significant differences in RLS across these strains on YPG condition [16]. Interestingly, a comparison of RLS phenotypes of these strains from YPG and YPD-CR revealed a significant positive correlation (R^2^ = 0.41, *P* = 6.18x 10-6) (**Supplementary Figure S3**) that strains responding to CR also respond to YPG positively or vice versa (**Figure 12, Supplementary File S1**). This data further confirms mitochondria associated with overlapping mechanisms mediating lifespan under glycerol and CR conditions.

**Figure 12:**
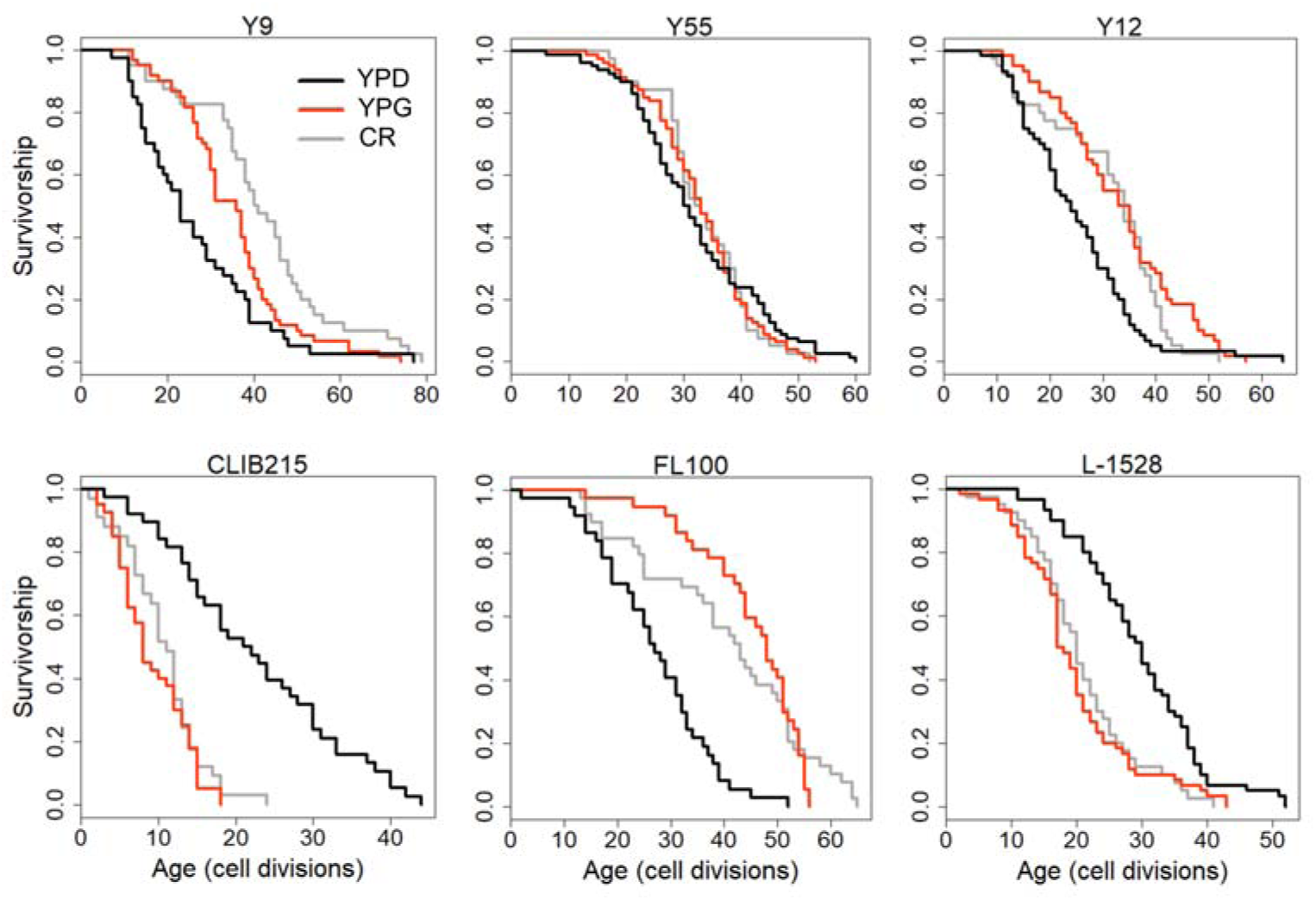
Examples of lifespan curves for the selected strains of *S. cerevisiae* analyzed under different metabolic conditions. RLS assays were conducted under control YPD (2% glucose, black lines), YPD-CR (0.05% glucose, gray lines), and YPG (3% glycerol, red lines) medium conditions. The raw data and statistical significance can be found in Supplementary File 1.

Next, to investigate the mitochondrial role in CR-mediated lifespan extension further, we eliminated mtDNA in two responding strains to isolate respiratory deficient cells [32,33]. Then, we analyzed their RLS under control and CR conditions. Elimination of mtDNA caused a decrease in median RLS in all three of them under 2% glucose condition (**Figure 13A, Supplementary File S1**). These results indicate mitochondria-specific metabolic adaptation in wild isolates since the elimination of mtDNA has shown mixed effect on median RLS in laboratory-adapted strains [33,34]. Next, we found that eliminating respiratory deficiency blocked the CR effect on lifespan extension (**Figure 13A**). Together, these data pinpoint that CR-mediated lifespan extension requires functional mitochondria in wild yeast isolates.

**Figure 13:**
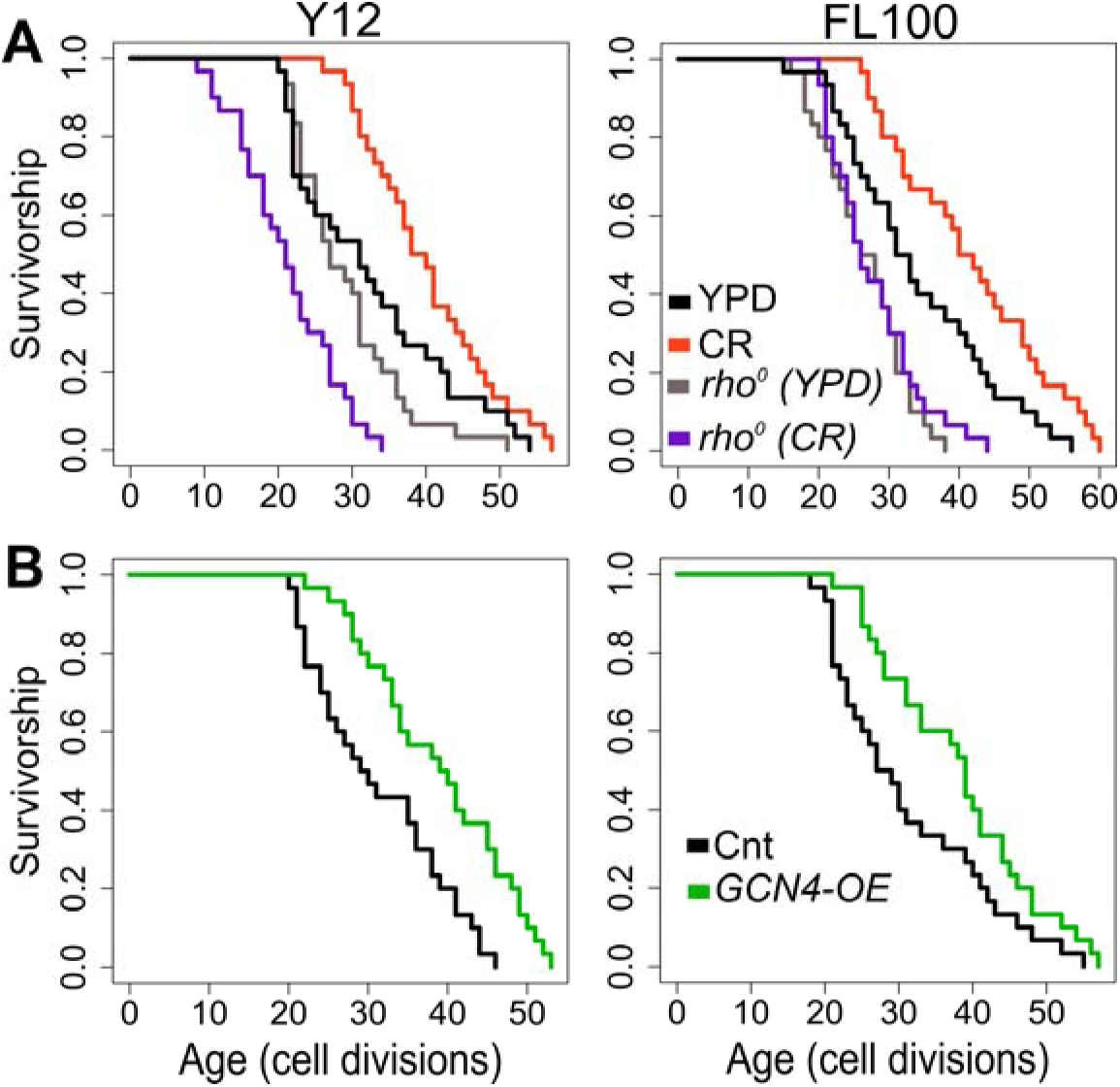
RLS effect of mitochondrial DNA (mtDNA) elimination and *GCN4* overexpression in selected responding strains. RLS phenotypes of Y12 (left panels) and FL100 (right panels) strains (**A**) rho0 isolates under high glucose condition (gray lines) and CR condition (purple lines) along with the RLS phenotypes of control parental strains under high glucose (YPD: black lines) and CR conditions (red lines) are also shown (**B**) Lifespan curves for control (black) and *GCN4* (green) overexpression. Lifespan data and the significance of lifespan changes can be found in Supplementary File 1.

Finally, we wanted to test whether increased *GCN4* expression alone can promote lifespan extension in selected CR responding strains (**Figure 13B**). Our transcriptome data showed that CR induces *GCN4* expression and alters *GCN4*-mediated pathways in CR responding strains (**Figure 12**). We further found that *GCN4* overexpression alone could induce lifespan extension (**Figure 13B, Supplementary File S1**). Our data represent unique adaptive changes involving *GCN4* function to mitochondria-mediated metabolic and molecular adaptations to the environment as the main factor for lifespan regulation under CR conditions.

## DISCUSSION

The initial studies reporting lifespan extension from reduced food intake without malnutrition, referred to as caloric restriction (CR) or dietary restriction (DR), were carried out more than a century ago in female rats [35]. Soon thereafter, similar observations were made in the water flea *Daphnia longispina* [36], in brook trout [37], and most definitively in the pioneering rat studies of McKay and colleagues [38]. Since then, CR has been repeatedly shown to induce lifespan extension and improve health outcomes across a broad range of evolutionarily distant organisms, including yeast, worms, flies, mice, and monkeys in laboratory settings [1–5]. Considerable research has been directed toward understanding CR-mediated lifespan regulation mechanisms. Consequently, several conserved genes and nutrition signaling pathways (e.g., mTOR) have been identified [39–41]. Currently, CR is considered one of the most promising dietary interventions for extending the lifespan of humans [10].

However, less attention has been paid to genotype-dependent responses to CR in the field [42,43]. There have been few reports exploring the effect of CR on the health and lifespan of genetically diverse flies [11,44] and mice were published [15,45, 46]. One commonly overlooked problem is most of the studies have been performed on laboratory-adapted model organisms, which have renewed our interest in understanding how the environment and interventions modulate lifespan diversity, leading to extended lifespan without significant reduction in fitness or fecundity [42, 46–49]. On the other hand, there have been a few reports from different model organisms, including yeast [50], housefly [51], and mice [52], that CR did not show significant lifespan effects or even reduced lifespan in some cases. Although some of these observations can be attributed to the methodologies [13] (e.g., the amount of restriction on the diet), these studies from closely related species indicated that genetic background is an important factor for CR response.

In this study, we tried to advance the understanding of the effect of genetic background on CR-mediated lifespan regulation and CR response by utilizing comparative genomics approaches using transcriptome signatures across highly diverse aging phenotypes of yeast isolates collected from different ecological niches [16–18]. Our RLS analyses showed that CR varies significantly within and between populations across different genotypes and species of budding yeast. While we observed that CR mediated positive response in some budding yeast species and strains of *S. cerevisiae*, most did not respond or decreased the lifespan, concluding that even among the closely-related species or strains of a single species, CR might not mediate lifespan extension commonly.

Next, a comparison of gene expression patterns between CR responding and non-responding strains revealed unique sets of genes and pathways and regulators of CR response. We found that responding wild strains uniquely adapted to increase mitochondrial translation coupled with oxidative phosphorylation and ATP biosynthesis under high glucose conditions. The laboratory-adapted budding yeast strain of *S. cerevisiae* mainly generates energy through glycolysis, and high glucose conditions suppress mitochondrial respiration. It is possible that niche-specific adaptations to the carbon source might be associated with increased mitochondrial function in these strains [16]. In addition, responding strains were characterized by decreased amino acid biosynthesis and increased cytosolic translation in comparison to both non-responding and negatively responding strains. In another observation, in addition to the decreased mitochondrial translation, non- and negatively responding strains are also characterized by respiratory chain complex IV assembly, inner mitochondrial membrane organization, cytochrome complex assembly, and ATP biosynthesis. This data suggests that under high glucose conditions (2%), negatively responding strains might have decreased mitochondrial capacity for energy derivation by oxidative phosphorylation, indicating adaptation of higher respiratory metabolisms under high glucose condition in responding strains. Overall, our data suggest mitochondrial function, cytosolic translation, and amino acid biosynthesis as the main factors for CR response type, and we further showed that two TFs, *HAP4* and *GCN4,* differentially regulate these pathways in responding strains.

There have been reports that focused on elucidating the *GCN4*-dependent lifespan regulation mainly in yeast and worm models *(ATF-4;* worm and mammalian functional orthologues of yeast *GCN4*) [53–58]. *GCN4* is a primary transcriptional activator of amino acid biosynthesis genes in yeast and regulates various stress resistance mechanisms [59–64]. Increased expression of *GCN4* has been shown to increase yeast and worm lifespan, and increased expression of *ATF4* was revealed to be a hallmark shared by many long-lived mice compared to those of normal-lived mice [65–66]. Increased *GCN4* expression was also found to decrease overall protein translation by regulating the expression of genes involved in translational machinery [56, 67]. Furthermore, long-lived yeast strains that lack non-essential ribosomal proteins have been characterized by an increased abundance of *GCN4,* and the increased RLS of these strains is mostly suppressed by the deletion of *GCN4* [56,68], indicating Gcn4-dependent longevity regulation in these strains. Interestingly, other studies also showed longevity promoting the function of Gcn4, independent of reduced global protein synthesis but through autophagic regulation in both yeast and mammalian cells [54,69]. Finally, consistent with our result of increased *GCN4* transcription under CR condition, the translational efficiency of Gcn4 was found to have significantly increased under CR [8,70,71], and it was also reported that Gcn4 is partially required for life span extension by CR [68].

Similarly, studies have demonstrated that the plasticity of mitochondrial function could be a potential target to promote healthy aging [72,73]. Mitochondria have an important role in a wide variety of metabolic and cellular processes, including energy production, amino acid synthesis, lipid metabolism, cell cycle regulation, apoptosis, autophagy, and signaling processes, and many of these processes are directly linked to lifespan regulation and aging [74,75]. However, mitochondrial function in aging and lifespan is more complex. For example, altered mitochondrial function is tightly linked to lifespan regulation, but underlying mechanisms remain unclear. The age-associated decline in mitochondrial function is associated with aging hallmarks and age-related diseases 76,77]. Modulation of mitochondria-related pathways through genetic, environmental, and pharmacological interventions has been shown to mediate longevity*-*promoting metabolic and molecular changes in different organisms, including yeast [17,78,79] *C. elegans* [80], and mice [81]. On the other hand, inhibition of mitochondrial respiration has also been associated with increased lifespan in various species [33, 82–85].

As a facultative anaerobe, yeast has been a valuable model for studying the role of mitochondrial function in aging. However, the continuous utilization of laboratory-adapted yeast has been the primary model in these studies and introduced some conflicting results. For example, the elimination of mtDNA (respiratory deficiency) has resulted in decreased [16] and increased [33] lifespan across different strains. In addition, although CR has been proposed to mediate its longevity effect through mitochondrial function in yeast [86], strains lacking mtDNA were also found to extend their lifespan under CR conditions [34]. Recently, we showed that the elimination of mtDNA decreases lifespan in many wild yeast isolates, indicating adapted metabolic changes in the laboratory environment, illustrating that the species has the potential genotype-specific lifespan traits regulated by particular genes in laboratory settings and might introduce artifact results [16]. In this study, we showed mitochondrial function is needed for CR-mediated lifespan extension in responding strains.

Although there have been numerous reports on understanding the mechanism of CR-mediated lifespan extension using the yeast model, our results represent the first comprehensive report on CR response type and associated molecular factors across the diverse lifespans of different genotypes. Our result identifies the possible interplay between Gcn4 and mitochondrial function together, mediating molecular and metabolic changes that ultimately exert a significant influence on the determination of longevity under CR conditions. Regulation of mitochondrial function and mitochondrial stress response, known as an integrated stress response (ISR) have been characterized in various organisms [87–90]. For example, inhibition of the functional copy of *ATF4* failed to up-regulate several mitochondrial enzymes and exhibited a reduction in ATP-dependent respiration in HeLa cells [88]. In the natural environment, *GCN4* might mediate signaling pathways connecting nutrient-sensing pathways to mitochondrial metabolic adaptation in response to nutrient availability to regulate phenotypic plasticity under varying stress environments. In fact, we showed that most of the up-regulated genes were previously characterized as a component of ISR [29]. Together, the *GCN4*-mediated ISR activation might be coupled to maintain mitochondrial and cellular homeostasis and organismal fitness under CR condition.

It should be noted here that we identified increased *GCN4* expression and decreased cytosolic translation in both groups. However, increased expression of mitochondrial genes was only observed in non-responsive strains under CR conditions. Responsive strains were characterized by higher mitochondrial activity and increased cytosolic translation under high glucose conditions compared to the non-responsive strains and CR did not cause further increases in mitochondrial function. The data suggest CR selectively acted only to decrease ribosomal biogenesis genes on responsive strains to extend lifespan. A similar observation was reported previously that knockdown of mitochondrial prohibitin *phb-2* induces the UPR^mt^ in both yeast and worms and shortens lifespan in both organisms. This effect of prohibitin deficiency can be suppressed by a reduction in cytoplasmic translation in yeast or by deletion of the S6 kinase homolog *rsks-1* in worms, which both increases the *GCN4/ATF4* function and attenuates the UPR^mt^ [7,91]. Although it needs further research, we argue that the absence of lifespan extension in non-responsive strains, even in the event of increased mitochondrial respiration, might be linked to the severely decreased cytosolic translation, indicating CR might cause imbalanced cytosolic protein homeostasis, which is tightly linked to mitochondrial translation efficiency and nuclear stress signaling [28,92].

In addition, we identified increased expression of methionine and other sulfur-containing amino acid biosynthesis genes regulated by *MET32* in non-responding strains under CR condition. Given that decreased methionine biosynthesis is required for CR-mediated lifespan extension, increased methionine biosynthesis might abrogate CR-mediated lifespan extension, as it was shown previously [71,93]. Further research is needed on the mechanisms that differentially promote increased expression of methionine biosynthetic genes in non-responding strains in comparison to the responding strains under CR condition. Another piece of evidence linking methionine metabolism to CR response emerged from our analysis of metabolomics data that increased abundance of n-formylmethione (fMet) under high glucose condition positively correlates with the rate of lifespan extension, promoted by CR. fMet is a derivate of methionine in which one of the hydrogens attached to the nitrogen in amino group is replaced by a formyl group and mainly compartmentalized in mitochondria, there it is used for initiation of mitochondrial protein synthesis [94,95]. Our data indicates that increased fMEt might be another key factor in coordinating the nutrient status and the mitochondrial translation linking CR-induced mitochondrial respiration. To our knowledge, there have been no reports showing fMet association with CR-mediated lifespan regulation and further research is needed whether fMet can be utilized as a predictive marker for CR response type and rate in higher eukaryotes.

Our comparison of transcriptome data revealed lower mitochondrial respiratory capacity under high glucose conditions for negatively responding strains. In addition, metabolomics data showed increased oxidized glutathione levels in them. The negatively responding strains may be suffering from oxidative stress, and switching the CR medium might activate mitochondrial respiration, further increasing the intracellular reactive oxygen species and thus decreasing lifespan.

Overall, our research has uncovered molecular determinants of lifespan plasticity in response to nutrition signaling that in the natural environment, it is employed to modify genotype and gene expression, arriving at different lifespans. However, our study only tested a single CR condition (0.05% glucose) that different levels of CR may interact with genotype in different ways. In particular, each genotype may have a different optimal (for longevity) level of nutrient availability, or the activity of metabolic pathways might differ significantly depending on genetic background under CR condition, as it was shown in mice previously [13,96,97]. Further understanding of how gene-environment interaction modulates genotype-dependent conserved molecular responses to various level of nutrition availability may open new therapeutic applications to slow aging and age-related diseases through diet, lifestyle, or pharmacological interventions.

## Materials and methods

### Yeast Strains and growth conditions

Diploid wild isolates of *S. cerevisiae* were obtained from the Sanger Institute [18], and the other budding yeast species were gifted by Mark Johnston [98]. We also reported detailed information and lifespan phenotypes of these strains analyzed under high glucose (2% glucose) and glycerol (3% glycerol) conditions [16,17]. For the expression of *GCN4*, we used a modified *p426ADH* plasmid by inserting a hygromycin (HYG) cassette along with its promoter and terminator at the *XbaI* restriction site [16]. We omitted the 5′UTR sequence of the *GCN4* as described previously [56]. Yeast transformation was performed using the standard lithium acetate method. Finally, to isolate *rho^0^* strains, cells were cultured in YPD medium, supplemented with 10 µg/ml and ethidium bromide (EtBr), and incubated at room temperature with agitation for approximately 24 hr. This procedure was repeated three times, and after the third growth cycle, cells were diluted (1:100) in water and plated on YPD to obtain single colonies. After that, several individual colonies were selected to test their growth ability on YPG (respiratory carbon source) plates. Colonies that were unable to grow on YPG were selected as *rho^0^*.

### Replicative Lifespan Analysis

RLS assay method and lifespan phenotypes of these strains on YEP (yeast extract, peptone) medium supplemented with 2% glucose or 3% glycerol was described in detail in our previous publications [16,17,99]. We modified the YEP plates by supplementing them with 0.05% glucose (YPD-CR) to determine the RLS phenotypes on the CR medium. For each natural isolate, at least 30 individual mother cells were analyzed. Each assay also included the BY4743 strain as a technical control. For RLS analysis of wild isolates harboring expression plasmids for *GCN4*, several individual colonies were picked up from selection medium (HYG) after transformation, and YPD medium supplemented with 200 µg/ml HYG was used for RLS determination of these cells. Survival analysis and Gompertz modeling were performed using the survival (https://cran.r-project.org/web/packages/survival/index.html) and flexsurv (https://cran.r-project.org/web/packages/flexsurv/index.html) packages in R, respectively.

### Comparison of gene expression and metabolomics signature associated with RLS phenotypes across strains

The RNAseq and metabolomics procedure and the data analyses for cells collected on YPD (2% glucose) were described previously [16]. We used the same procedure to collect cells and perform RNAseq analyses for Saccharomyces species and for the cells collected don YPD-CR (0.05% glucose) medium. Data analyses were also done by using similar methodologies for consistency.

To assess the impact of experimental treatments on gene expression, we conducted a Principal Component Analysis (PCA) utilizing the “FactoMineR” package in R [100]. The PLS-DA analysis was performed by using the function ‘opls’ in R package ‘ropls’ with default parameters (https://bioconductor.org/packages/devel/bioc/vignettes/ropls/inst/doc/ropls-vignette.html#43_Partial_least-squares:_PLS_and_PLS-DA). Additionally, we computed the Spearman correlation coefficient between samples based on their gene expression values. The distance matrix among samples was derived from the 1−correlation coefficient and employed the “hclust” function in R for hierarchical clustering using the “ward.D2” method. For the differential expression analysis, we utilized the “EdgeR” package in R [101] to perform comparisons between positively responding versus non- and negatively responding groups. Briefly, we initially applied the “calcNormFactors” function to calculate the normalization factor for the raw counts of gene expressions by using the value of the total expression of samples. Subsequently, we used the “voomWithQualityWeights” function to estimate the weight of each sample and transformed the expression matrix using voom. Given that there are multiple strains within each group, with each strain having three biological replicates, treating biological repetitions as independent observations would introduce noise into the results. To address this, we employed the “duplicateCorrelation” function to calculate the correlation among biological replicates. We then reapplied the “voomWithQualityWeights” function to re-estimate the sample weights and transform the expression matrix, considering the correlation between biological replicates. Finally, we utilized the “lmFit” function to identify the significant differential expressed gene by setting each strain as a block and incorporating the correlation between biological replicates. The adjustment of the *P* value was performed using the Benjamini-Hochberg (BH) method. Gene met adjusted *P* value < 0.05 and | log2 (Fold change) | > 0.5 as the significant genes. To identify endophenotypes (transcripts and metabolites) correlating with CR-mediated lifespan variation across wild isolates, we performed phylogenetic regression using the generalized least squares method, as we described previously [16,17].

### Functional enrichment and transcription factor network analysis

We performed gene enrichment analysis using R packages “clusterProfiler” [102]. We selected the items contained in the Gene ontology (GO) and the KEGG databases to analyze up- and down-regulated genes for each group. The Benjamini-Hochberg (BH) method was used for the *P* value adjustment, and the terms with adjusted *P* value < 0.05 were selected. For the KEGG database-based pathway enrichments, terms with adjusted *P* values < 0.5 were considered. To identify TFs associated with DEGs identified under CR conditions, we utilized the yeastrack portal [103], which provides a set of queries to predict transcription regulation networks in yeast from various -omics data.

## Author contributions

SM, ML, RT and AK performed the experiments. SM, ML, VAG, WZ, XZ and AK analyzed the data. SM, ML, VAG, WZ, RT XZ, MK and AK drafted and revised the manuscript. All authors approved the submitted version of the manuscript.

## Acknowledgments

We would like to acknowledge the generous funding provided by the NIA/NIH (1K01AG060040) and NIGMS/NIH (1R35GM150858-01) to AK. Studies performed by M. K. were funded by the University of Washington Nathan Shock Center of Excellence in the Basic Biology pilot award to A. K. (P30AG013280). The funders had no role in study design, data collection and interpretation, or the decision to submit the work for publication.

## Data availability

All data generated or analyzed during this study are included in the manuscript and supporting files. RNA-seq data for the cells collected under high glucose conditions were previously deposited to the NCBI Gene Expression Omnibus (GEO) with accession number GSE188294 and RNA-seq data for the cells collected under low glucose (CR) conditions are deposited to the NCBI Gene Expression Omnibus (GEO) with accession number XX.

## Supplementary Figures

**Figure S1:**
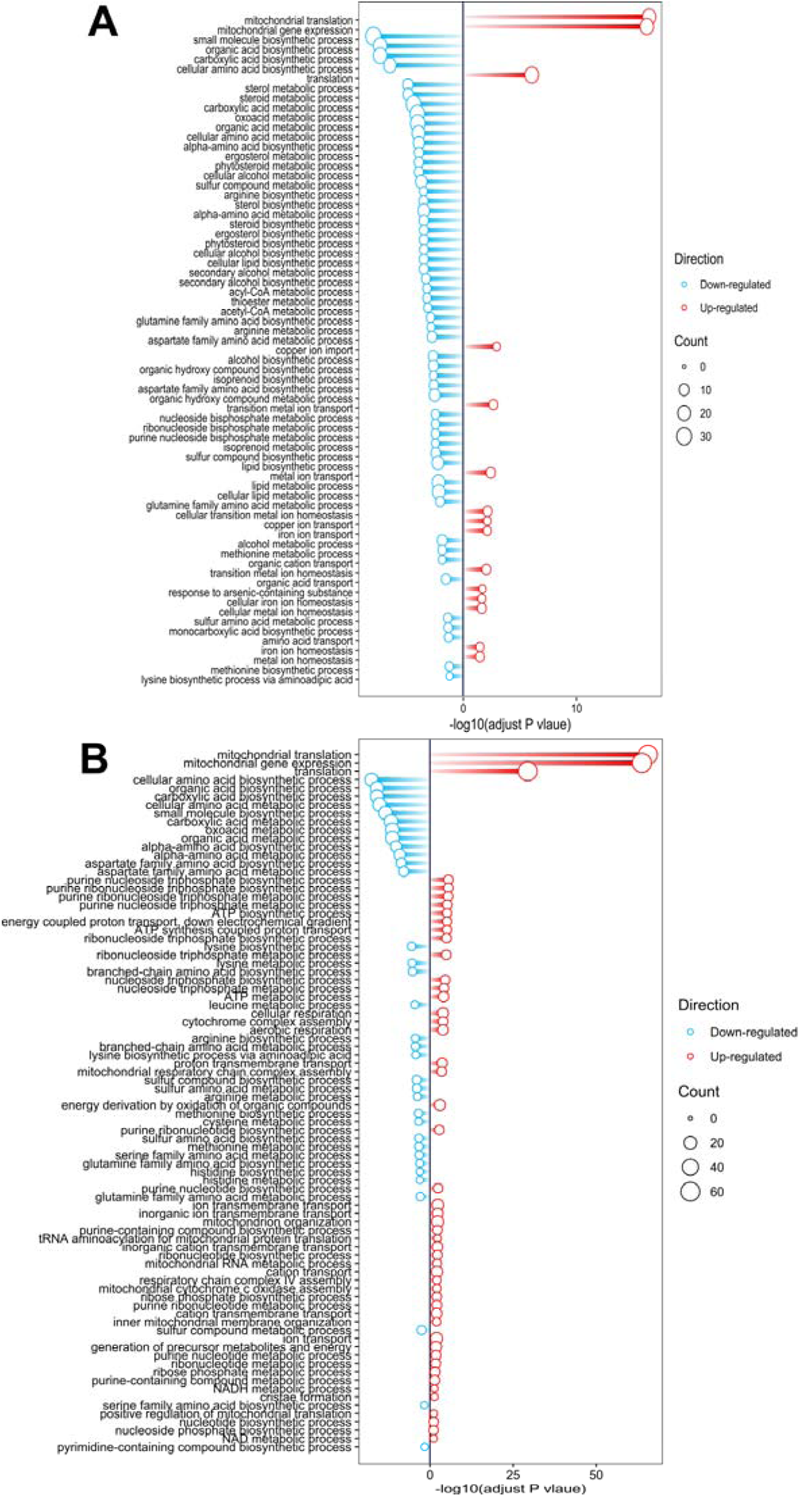
GO term enrichment analyses: The GO term (Molecular Function) analyses show significantly enriched terms for (**A**) positively responding versus non-responding and (**B**) positively responding versus negatively responding strains.

**Figure S2.**
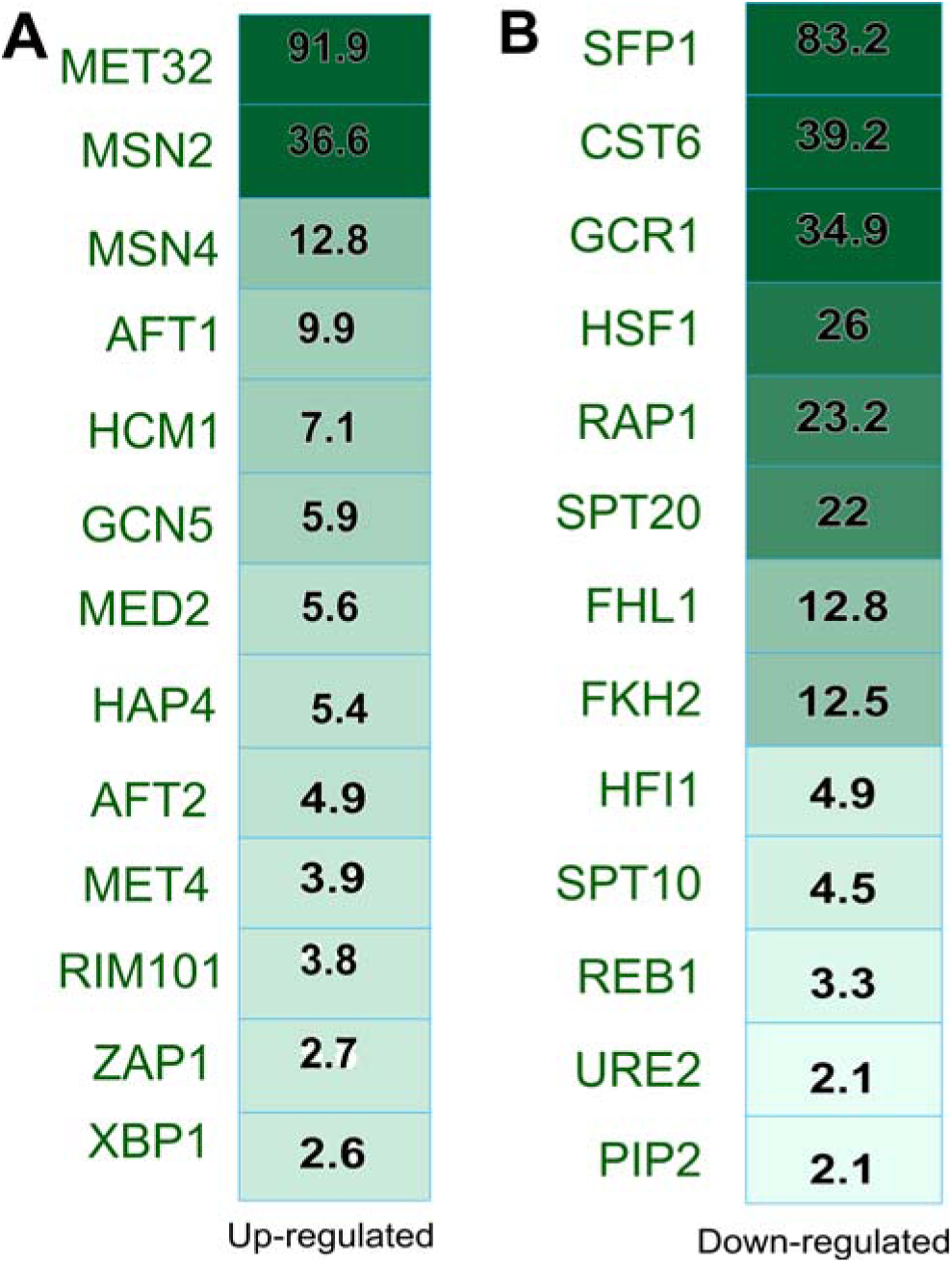
Transcription factor enrichment analyses for non-responding strains analyzed under CR condition. Significantly enriched TFs for (**A**) up- and (**B**) down-regulated DEGs obtained under CR conditions for non-responding strains. The color index and the number show the significance of the enrichment score (-log_10_, adjusted *P*-value).

**Figure S3.**
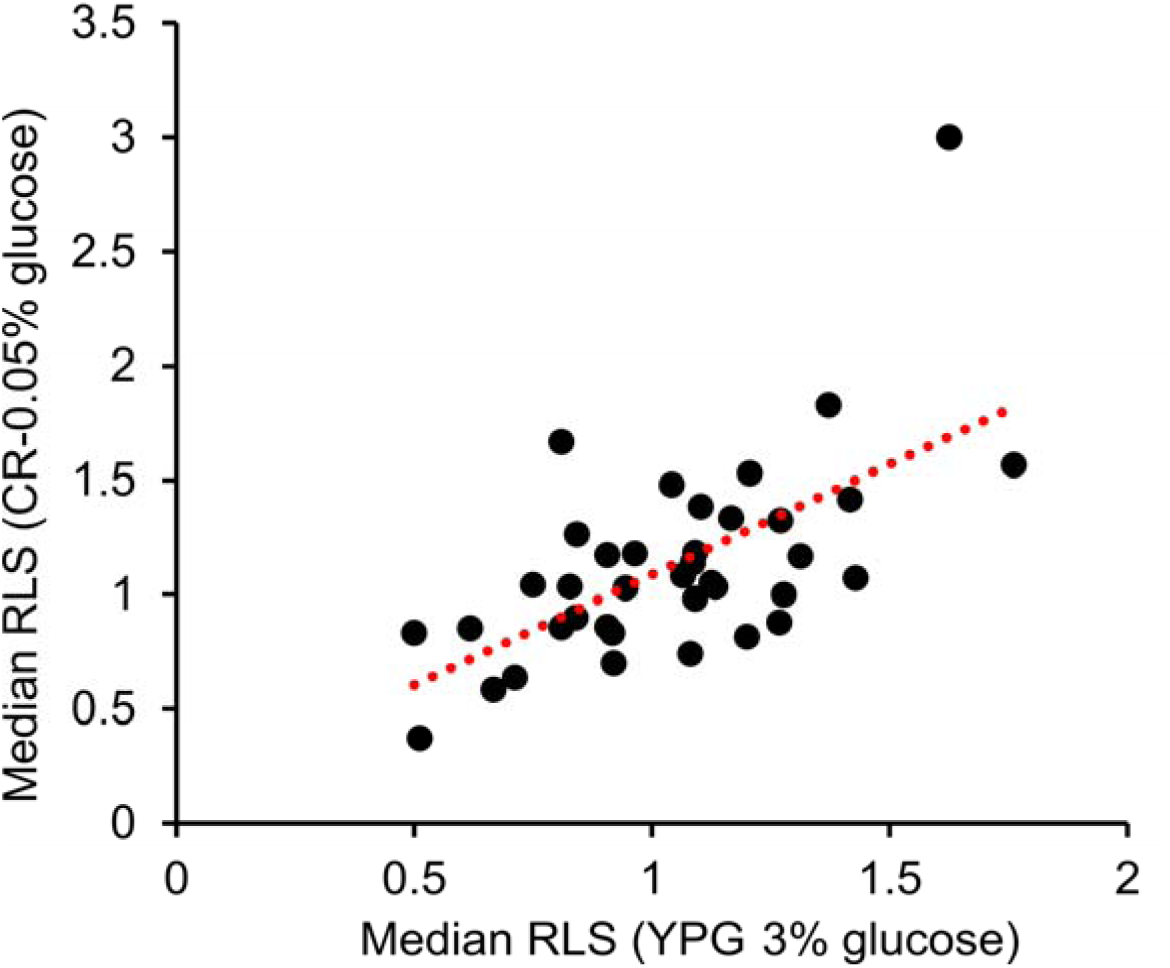
A positive correlation of median RLS phenotype between YPG and CR medium was observed. A significant correlation was observed between median RLS in YPG and median RLS in CR, wherein glycerol-induced CR-induced respiration approximately equally extends the RLS of those strains we tested (R^2^ = 0.62, Padj value = 2.29x10-4).

## Supplementary Files

**Supplementary file 1:** The file includes the strain list used in this study and their replicative lifespan analyzed under YPD, YPG, and CR conditions, along with the statistical significance of lifespan differences.

**Supplementary File 2:** Differentially expressed genes and metabolite abundances resulted from comparison of responding (POS) versus non-responding (NON) and negatively (NEG) responding comparisons. These -omics data were obtained under YPD conditions. The file also contains results from the regression analysis for the genes and metabolites whose transcript level/abundances correlate with CR-lifespan. The file also includes results from the GO term enrichment analysis associated with Figures 9 and 10.

**Supplementary File 3:** The file includes raw and normalized counts for RNA-seq data for selected responding and non-responding strains. Differentially expressed genes for each group resulting from comparison to the control (YPD) are also included.

## References

1. Anderson, R. M. & Weindruch, R. The caloric restriction paradigm: Implications for healthy human aging. American Journal of Human Biology 24, 101–106 (2012).

2. López-Lluch, G. & Navas, P. Calorie restriction as an intervention in ageing. Journal of Physiology 594, 2043–2060 (2016).

3. Hoshino, S., Kobayashi, M. & Higami, Y. Mechanisms of the anti-aging and prolongevity effects of caloric restriction: evidence from studies of genetically modified animals. Aging 10, 2243–2251 (2018).

4. Sinclair, D. A. Toward a unified theory of caloric restriction and longevity regulation. Mech Ageing Dev 126, 987–1002 (2005).

5. Ungvari, Z., Parrado-Fernandez, C., Csiszar, A. & De Cabo, R. Mechanisms underlying caloric restriction and lifespan regulation: Implications for vascular aging. Circulation Research vol. 102, 519–529 (2008).

6. Lin, S.-J. et al. Calorie restriction extends Saccharomyces cerevisiae lifespan by increasing respiration. Nature 418, 344–348 (2002).

7. Schleit, J. et al. Molecular mechanisms underlying genotype-dependent responses to dietary restriction. Aging Cell 12, 1050–1061 (2013).

8. Zou, K. et al. Life span extension by glucose restriction is abrogated by methionine supplementation: Cross-talk between glucose and methionine and implication of methionine as a key regulator of life span. Sci. Adv 6(2):eaba1306 (2020).

9. Steffen, K. K., Kennedy, B. K. & Kaeberlein, M. Measuring replicative life span in the budding yeast. J Vis Exp 28, 1209 (2009)

10. Lee, M. B., Hill, C. M., Bitto, A. & Kaeberlein, M. Antiaging diets: Separating fact from fiction. Science 374(6570):eabe7365 (2021).

11. Jin, K. et al. Genetic and metabolomic architecture of variation in diet restriction-mediated lifespan extension in Drosophila. PLoS Genet 16(7):e1008835 (2020).

12. McCracken, A. W., Buckle, E. & Simons, M. J. P. The relationship between longevity and diet is genotype dependent and sensitive to desiccation in Drosophila melanogaster. Journal of Experimental Biology 223(Pt 23):jeb230185 (2020).

13. Mitchell, S. J. et al. Effects of Sex, Strain, and Energy Intake on Hallmarks of Aging in Mice. Cell Metab 23, 1093–1112 (2016).

14. Stastna, J. J., Snoek, L. B., Kammenga, J. E. & Harvey, S. C. Genotype-dependent lifespan effects in peptone deprived Caenorhabditis elegans. Sci Rep 5, 16259 (2015).

15. Liao, C. Y., Rikke, B. A., Johnson, T. E., Diaz, V. & Nelson, J. F. Genetic variation in the murine lifespan response to dietary restriction: From life extension to life shortening. Aging Cell 9, 92–95 (2010).

16. Kaya, A. et al. Evolution of natural lifespan variation and molecular strategies of extended lifespan in yeast. Elife 10:e64860 (2021).

17. Kaya, A. et al. Defining molecular basis for longevity traits in natural yeast isolates. NPJ Aging Mech Dis 1, 15001 (2015).

18. Liti, G., et al. Population Genomics of Domestic and Wild Yeasts. Nature vol. 458, 337–341 (2009).

19. Gilad, Y., Oshlack, A. & Rifkin, S. A. Natural selection on gene expression. Trends in Genetics 22, 456–461 (2006).

20. Vu, V. et al. Natural Variation in Gene Expression Modulates the Severity of Mutant Phenotypes. Cell 162, 391–402 (2015).

21. Liu, W. et al. LargeLscale across species transcriptomic analysis identifies genetic selection signatures associated with longevity in mammals. EMBO J 42(17):e112740. (2023).

22. Whitaker, R. et al. Dietary switch reveals fast coordinated gene expression changes in Drosophila melanogaster. Aging 6, 355–368 (2014).

23. Brawand, D. et al. The evolution of gene expression levels in mammalian organs. Nature 478, 343–348 (2011).

24. Laye, M. J., Tran, V., Jones, D. P., Kapahi, P. & Promislow, D. E. L. The effects of age and dietary restriction on the tissue-specific metabolome of Drosophila. Aging Cell 14, 797–808 (2015).

25. Ma, S. et al. Organization of the Mammalian Metabolome according to Organ Function, Lineage Specialization, and Longevity. Cell Metab 22, 332–343 (2015).

26. Cheng, S. et al. Distinct metabolomic signatures are associated with longevity in humans. Nat Commun 6, 6791 (2015).

27. McCarthy, D. J., Chen, Y. & Smyth, G. K. Differential expression analysis of multifactor RNA-Seq experiments with respect to biological variation. Nucleic Acids Res 40, 4288–4297 (2012).

28. Molenaars, M. et al. A Conserved Mito-Cytosolic Translational Balance Links Two Longevity Pathways. Cell Metab 31, 549–563.e7 (2020).

29. Gasch, A. P., et al. Genomic Expression Programs in the Response of Yeast Cells to Environmental Changes D. Molecular Biology of the Cell 12, 4241– 4257 (2000).

30. Burtner, C. R., Murakami, C. J., Kennedy, B. K. & Kaeberlein, M. A molecular mechanism of chronological aging in yeast. Cell Cycle 8, 1256–1270 (2009).

31. Kaeberlein, M. et al. Regulation of Yeast Replicative Life Span by TOR and Sch9 in Response to Nutrients. Science 310, 1193–1196 (2005).

32. Kaya, A. et al. Adaptive aneuploidy protects against thiol peroxidase deficiency by increasing respiration via key mitochondrial proteins. Proc Natl Acad Sci U S A 112, 10685–10690 (2015).

33. Woo, D. K. & Poyton, R. O. The absence of a mitochondrial genome in rho0 yeast cells extends lifespan independently of retrograde regulation. Exp Gerontol 44, 390–397 (2009).

34. Kaeberlein, M. et al. Increased life span due to calorie restriction in respiratory-deficient yeast. PLoS Genet 3:e33 (2005).

35. Osborne, T. B., Mendel, L. B. & Ferry, E. L. The Effect of Retardation of Growth Upon the Breeding Period and Duration of Life of Rats. Science 45, 294–295 (1917).

36. Ingle, L., Wood, T. R. & Banta, A. M. A study of longevity, growth, reproduction and heart rate in Daphnia longispina as influenced by limitations in quantity of food. Journal of Experimental Zoology 76, 325–352 (1937).

37. Titcomb, J. W., Cobb, E. W., Crowell, M. F. & McCay, C. M. The nutritional requirements and growth rates of brook trout. Trans Am Fish Soc 58, 205–231 (1928).

38. Mccay, C. M., Crowell, M. P. & Maynakd, L. A. The effect of retarded growth upon the length of life span and upon the ultimate body size. Nutrition 5; 155–171 (1989).

39. Green, C. L., Lamming, D. W. & Fontana, L. Molecular mechanisms of dietary restriction promoting health and longevity. Nat Rev Mol Cell Biol 23, 56–73 (2022).

40. Komatsu, T. et al. Mechanisms of calorie restriction: A review of genes required for the life-extending and tumor-inhibiting effects of calorie restriction. Nutrients 11, 3068 (2019).

41. Wu, Q., Gao, Z. J., Yu, X. & Wang, P. Dietary regulation in health and disease. Signal Transduct Target Ther 7, 252 (2022).

42. Selman, C. & Swindell, W. R. Putting a strain on diversity. EMBO J 37(22):e100862 (2018).

43. Liao, C. Y., Johnson, T. E. & Nelson, J. F. Genetic variation in responses to dietary restriction - An unbiased tool for hypothesis testing. Exp Gerontol 48, 1025–1029 (2013).

44. Wilson, K. A. et al. GWAS for Lifespan and Decline in Climbing Ability in Flies upon Dietary Restriction Reveal decima as a Mediator of Insulin-like Peptide Production. Current Biology 30, 2749–2760 (2020).

45. 45. Di Francesco, A. et al. Regulators of health and lifespan extension in genetically diverse mice on dietary restriction. bioRxiv 2023.11.28.568901; doi: 10.1101/2023.11.28.568901.

46. Chen, J., et al. A Demographic Analysis of the Fitness Cost of Extended Longevity in Caenorhabditis Elegans. J Gerontol A Biol Sci Med Sci. 62, 126–35 (2007).

47. Jenkins, N. L., McColl, G. & Lithgow, G. J. Fitness cost of extended lifespan in Caenorhabditis elegans. Proc Biol Sci 271, 2523–2526 (2004).

48. Maklakov, A. A. & Chapman, T. Evolution of ageing as a tangle of trade-offs: Energy versus function. Proc R Soc B 286, 20191604 (2019).

49. Ramani, A. K. et al. The majority of animal genes are required for wild-type fitness. Cell 148, 792–802 (2012).

50. Huberts, D. H. E. W. et al. Calorie restriction does not elicit a robust extension of replicative lifespan in Saccharomyces cerevisiae. Proc Natl Acad Sci U S A 111, 11727–11731 (2014).

51. Cooper, T. M., Mockett, R. J., Sohal, B. H., Sohal, R. S. & Orr, W. C. Effect of caloric restriction on life span of the housefly, Musca domestica . The FASEB Journal 18, 1591–1593 (2004).

52. Forster, M. J., Morris, P. & Sohal, R. S. Genotype and age influence the effect of caloric intake on mortality in mice. The FASEB journalU**17**, 690–692 (2003).

53. Anderson, N. S. & Haynes, C. M. Folding the Mitochondrial UPR into the Integrated Stress Response. Trends Cell Biol 6, 428–439 (2020).

54. Hu, Z., et al. Ssd1 and Gcn2 suppress global translation efficiency in replicatively aged yeast while their activation extends lifespan. Elife 7:e35551 (2018).

55. Mariner, B. L., Felker, D. P., Cantergiani, R. J., Peterson, J. & McCormick, M. A. Multiomics of GCN4-Dependent Replicative Lifespan Extension Models Reveals Gcn4 as a Regulator of Protein Turnover in Yeast. Int J Mol Sci 24, 16163 (2023).

56. Mittal, N. et al. The Gcn4 transcription factor reduces protein synthesis capacity and extends yeast lifespan. Nat Commun 8, 457 (2017).

57. Statzer, C. et al. ATF-4 and hydrogen sulfide signalling mediate longevity in response to inhibition of translation or mTORC1. Nat Commun 13, 967 (2022).

58. Vasudevan, D. et al. Translational induction of ATF4 during integrated stress response requires noncanonical initiation factors eIF2D and DENR. Nat Commun 11, 4677 (2020).

59. Jia, M. H., et al. Global Expression Profiling of Yeast Treated with an Inhibitor of Amino Acid Biosynthesis, Sulfometuron Methyl. Physiol Genomics 3, 83–92 (2000).

60. Kreß, J. K. C. et al. The integrated stress response effector ATF4 is an obligatory metabolic activator of NRF2. Cell Rep 42, 112724 (2023).

61. Mascarenhas, C. et al. Gcn4 is required for the response to peroxide stress in the yeast Saccharomyces cerevisiae. Mol Biol Cell 19, 2995–3007 (2008).

62. Natarajan, K. et al. Transcriptional Profiling Shows that Gcn4p Is a Master Regulator of Gene Expression during Amino Acid Starvation in Yeast. Mol Cell Biol 21, 4347–4368 (2001).

63. Patil, C. K., Li, H. & Walter, P. Gcn4p and novel upstream activating sequences regulate targets of the unfolded protein response. PLoS Biol 8:E246 (2004).

64. Wortel, I. M. N., van der Meer, L. T., Kilberg, M. S. & van Leeuwen, F. N. Surviving Stress: Modulation of ATF4-Mediated Stress Responses in Normal and Malignant Cells. Trends Endocrinol Metab 28, 794–806 (2017).

65. Li, W. & Miller, R. A. Elevated ATF4 function in fibroblasts and liver of slow-aging mutant mice. Gerontol A Biol Sci Med Sci 70, 263–272 (2015).

66. Li, W., Li, X. & Miller, R. A. ATF4 activity: A common feature shared by many kinds of slow-aging mice. Aging Cell 13, 1012–1018 (2014).

67. Hinnebusch, A. G. Translational regulation of GCN4 and the general amino acid control of yeast. Annu Rev Microbiol 59, 407–450 (2005).

68. Steffen, K. K. et al. Yeast Life Span Extension by Depletion of 60S Ribosomal Subunits Is Mediated by Gcn4. Cell 133, 292–302 (2008).

69. Mariner, B. L., Rodriguez, A. S., Heath, O. C. & McCormick, M. A. Induction of proteasomal activity in mammalian cells by lifespan-extending tRNA synthetase inhibitors. Geroscience 46, 1755–1773 (2024).

70. Yang, R., Wek, S. A. & Wek, R. C. Glucose limitation induces GCN4 translation by activation of Gcn2 protein kinase. Mol Cell Biol 20, 2706–2717 (2000).

71. Mehta, R., Chandler-Brown, D., Ramos, F.J., Shamieh L.S. & Kaeberlein, M. Regulation of mRNA translation as a conserved mechanism of longevity control. Advances in Experimental Medicine and Biology. 694, 14–29 (2010).

72. Lima, T., Li, T. Y., Mottis, A. & Auwerx, J. Pleiotropic effects of mitochondria in aging. Nat Aging 2, 199–213 (2022).

73. Phua, C. Z. J. et al. Genetic perturbation of mitochondrial function reveals functional role for specific mitonuclear genes, metabolites, and pathways that regulate lifespan. Geroscience 45, 2161–2178 (2023).

74. Goodell, M. A. & Rando, T. A. Stem cells and healthy aging. Science 350, 1199–1204 (2015).

75. Jang, J. Y., Blum, A., Liu, J. & Finkel, T. The role of mitochondria in aging. J Clin Invest 128, 3662–3670 (2018).

76. Lenaz, G., et al. Mitochondrial Complex I Defects in Aging. Mol Cell Biochem 174, 329–333 (1997).

77. Pawley, J. The Case for Low Voltage High Resolution Scanning Microscopy of Biological Samples. Scanning Microsc Suppl 3, 163–78 (1989).

78. Bonawitz, N. D., Chatenay-Lapointe, M., Pan, Y. & Shadel, G. S. Reduced TOR Signaling Extends Chronological Life Span via Increased Respiration and Upregulation of Mitochondrial Gene Expression. Cell Metab 5, 265–277 (2007).

79. Ocampo, A., Liu, J., Schroeder, E. A., Shadel, G. S. & Barrientos, A. Mitochondrial respiratory thresholds regulate yeast chronological life span and its extension by caloric restriction. Cell Metab 16, 55–67 (2012).

80. Schulz, T. J. et al. Glucose Restriction Extends Caenorhabditis elegans Life Span by Inducing Mitochondrial Respiration and Increasing Oxidative Stress. Cell Metab 6, 280–293 (2007).

81. Anderson, R. M. & Weindruch, R. Metabolic reprogramming, caloric restriction and aging. Trends Endocrinol Metab 21, 134–141 (2010).

82. Copeland, J. M. et al. Extension of Drosophila Life Span by RNAi of the Mitochondrial Respiratory Chain. Curr Biol 19, 1591–1598 (2009).

83. Liu, X. et al. Evolutionary conservation of the clk-1-dependent mechanism of longevity: Loss of mclk1 increases cellular fitness and lifespan in mice. Genes Dev 19, 2424–2434 (2005).

84. Van Raamsdonk, J. M. et al. Decreased energy metabolism extends life span in Caenorhabditis elegans without reducing oxidative damage. Genetics 185, 559–571 (2010).

85. Lee, S. J., Hwang, A. B. & Kenyon, C. Inhibition of respiration extends C. elegans life span via reactive oxygen species that increase HIF-1 activity. Curr Biol 20, 2131–2136 (2010).

86. Lin, S.-J. et al. Calorie restriction extends Saccharomyces cerevisiae lifespan by increasing respiration. Nature 418, 344–348 (2002).

87. PakosLZebrucka, K. et al. The integrated stress response. EMBO Rep 17, 1374–1395 (2016).

88. Quirós, P. M. et al. Multi-omics analysis identifies ATF4 as a key regulator of the mitochondrial stress response in mammals. J Cell Biol 216, 2027–2045 (2017).

89. Sasaki, K. et al. Mitochondrial translation inhibition triggers ATF4 activation, leading to integrated stress response but not to mitochondrial unfolded protein response. Biosci Rep 40(11):BSR20201289 (2020).

90. Zid, B. M. et al. 4E-BP Extends Lifespan upon Dietary Restriction by Enhancing Mitochondrial Activity in Drosophila. Cell 139, 149–160 (2009).

91. Artal-Sanz, M. & Tavernarakis, N. Prohibitin couples diapause signalling to mitochondrial metabolism during ageing in C. elegans. Nature 461, 793–797 (2009).

92. Suhm, T. et al. Mitochondrial Translation Efficiency Controls Cytoplasmic Protein Homeostasis. Cell Metab 27, 1309–1322.e6 (2018).

93. Lee, B. C., Kaya, A. & Gladyshev, V. N. Methionine restriction and life span control. Ann N Y Acad Sci 1363, 116–124 (2016).

94. Cai, N. et al. Mitochondrial DNA variants modulate N-formylmethionine, proteostasis and risk of late-onset human diseases. Nat Med 27, 1564–1575 (2021).

95. Lee, C. S., Kim, D. & Hwang, C. S. Where Does N-Formylmethionine Come from? What for? Where Is It Going? What is the origin of N-formylmethionine in eukaryotic cells? Mol Cells 45, 109–111 (2022).

96. Mulvey L. et al. Strain-specific metabolic responses to long-term caloric restriction in female ILSXISS recombinant inbred mice. Mol Cell Endocrinol. 535,111376 (2021).

97. Wilkie S.E. et al. Strain-specificity in the hydrogen sulphide signalling network following dietary restriction in recombinant inbred mice. Geroscience 42, 801–812 (2020).

98. Cliften, P. F. et al. Surveying Saccharomyces genomes to identify functional elements by comparative DNA sequence analysis. Genome Res 11, 1175–1186 (2001).

99. Oz, N. et al. Evidence that conserved essential genes are enriched for pro-longevity factors. Geroscience 44, 1995–2006 (2022).

100. Lê, S., Josse, J., Rennes, A. & Husson, F. FactoMineR: An R Package for Multivariate Analysis. JSS Journal of Statistical Software 25,1–18 2008.

101. Robinson, M. D., McCarthy, D. J. & Smyth, G. K. edgeR: A Bioconductor package for differential expression analysis of digital gene expression data. Bioinformatics 26, 139–140 (2009).

102. Yu, G., Wang, L. G., Han, Y. & He, Q. Y. ClusterProfiler: An R package for comparing biological themes among gene clusters. OMICS 16, 284–287 (2012).

103. Teixeira, M. C. et al. YEASTRACT+: a portal for the exploitation of global transcription regulation and metabolic model data in yeast biotechnology and pathogenesis. Nucleic Acids Res 51, D785–D791 (2023).

